# On-chip engineered human lymphatic microvasculature for physio-/pathological transport phenomena studies

**DOI:** 10.1101/2022.03.06.483122

**Authors:** Jean C. Serrano, Mark R. Gillrie, Ran Li, Sarah H. Ishamuddin, Roger D. Kamm

## Abstract

The human vasculature constitutes an integral part of fluid, protein and cellular transport throughout a variety of physiological processes and pathological events. While the blood vascular system has been the topic of numerous studies in connection to its role in physio-/pathological transport phenomena, our secondary vascular system, the lymphatics, has yet to gain similar attention, in part due to a lack of adequate models to study its biological function. Despite their considerable value, animal models limit the ability to perform parametric studies, whereas current *in vitro* systems are lacking in physiological mimicry. Here, a microfluidic-based approach is developed that allows for precise control over the transport of growth factors and interstitial fluid flow, which we leverage to recapitulate the *in vivo* growth of lymphatic capillaries. Using this approach, physiological tissue functionality is validated by characterizing the drainage rate of extracellular solutes and proteins. Finally, lymphatic-immune interactions are studied to affirm inflammation-driven responses by the lymphatics, which recruit immune cells via chemotactic signals, similarly to *in vivo*, pathological events. Results demonstrate the utility of this platform to study lymphatic biology and disease, as well as use as a screening assay to predict lymphatic absorption of therapeutic biologics.

## 1. Introduction

Most human tissues contain a secondary vascular system known as the lymphatics comparable in complexity to the blood vasculature. Both systems serve as an elaborate, hierarchal network of vessels lined by endothelial cells that serve as conduits for fluid, protein and cellular transport [1]. However, in contrast to the blood vascular system where fluid is continuously recirculating through different tissues, the lymphatic system operates as a one-way transport pathway that collects fluid, proteins and cells from the interstitial space of tissues and returns them to the systemic circulation [2]. Under this mechanism, lymphatics contribute to fluid and osmotic pressure homeostasis in tissues. In addition, the lymphatic system acts as an immune checkpoint by transporting antigen and antigen-presenting immune cells from the interstitial tissue to the lymph node, where resident immune cells respond to localized or systemic inflammation and infections [3].

As critical as the lymphatics are for physiological fluid, protein and cellular transport, research towards understanding their biological function/interactions and pathological alterations has been severely lacking, especially when compared to the vast literature on blood vasculature [4]. One of the overriding challenges in lymphatics-focused studies has been the lack of experimental models that facilitate studies to interrogate biological mechanisms implicated in lymphatic development, physiology and disease. For years, animal models served as the gold standard to evaluate the biological function of lymphatics, as well as its implications in pathological events such as inflammation, pathogen response and cancer progression [5–7]. Despite fully recapitulating physiological responses, animal studies offer limited control over local environmental cues, present barriers to isolate and describe the direct and indirect systemic effects of the modulated parameters, and their findings are often difficult to relate to corresponding phenomena in humans due to species differences. To mitigate these limitations, numerous groups have implemented *in vitro* models to perform reductionist studies on lymphatic development and pathogenesis with the ability to isolate the individual contribution of regulated cues in the cellular microenvironment [8–11]. However, the simplicity of such models often leads to a lack of physiological relevance, limiting their applicability to study *in vivo* events.

Given these limitations, there is a need for next generation platforms that more fully recapitulate the complexities of the *in vivo* microenvironment, while leveraging tight control over the biological interactions under study. This has driven development of microfluidic technologies that address these needs by the culture of human-sourced cells in three-dimensional (3D) environments that mimic tissue architecture with precise manipulation of mass transport throughout the cellular microenvironment [12–14]. However, studies implementing microfluidic systems to generate lymphatic vasculature are exceedingly scarce. Apart from simple monolayer systems that lack anatomical resemblance, fewer than a dozen published works have attempted to recreate the native lymphatic vasculature on-chip [14–16]. Most of these studies implement a single lymphatic capillary on-chip system to study solute permeability, and paracrine signaling/conditioning between lymphatic and fibroblast/tumor cells [17–20]. While such studies have provided fundamental insight into the biology of lymphatics, these platforms have limited utility to study lymphatic tissue function given their simple, single-capillary structure which fails to recapitulate the branching hierarchical structure found *in vivo*.

As with most tissues and organs, the adequate patterning of lymphatic vascular structures arises from the precise spatiotemporal control of biomolecules and biomechanical stimuli which guide migration, growth and remodeling during development [21]. By emulating the biological transport of these signals, on-chip systems can exploit vascular self-assembly for tissue engineering lymphatic microvasculature that inherently captures its complex, native network structure [22]. Interestingly, mass transport not only plays a critical role in shaping the distribution of factors that give rise to these vascular structures, but also constitutes the fundamental phenomena by which lymphatic drainage, and lymphatic-immune biochemical signaling occurs [23]. However, to the best of our knowledge, no published work has yet utilized the advantages of precisely delivering biological factors, in a microfluidic-based platform, to reconstitute lymphatic vascular formation and physiological tissue-level function.

In this work, we describe a microfluidic approach to generate functional lymphatic microvascular networks in a 3D hydrogel compartment by leveraging the capability to control the spatio-temporal delivery of solutes, protein and human cells. To achieve this, we first screened for the optimal balance of growth factors, extracellular matrix composition and interstitial fluid flow that would induce controlled-levels of angiogenic sprouting by lymphatic endothelial cells. After validating the *in vivo*-like morphology of our engineered lymphatics, we quantified their clearance function obtaining results for solute drainage rates approaching *in vivo* measurements. We also analyzed the underlying transport phenomena, elucidating the importance of a 3D geometry and the lymphatic endothelium to recapitulate physiological drainage. Finally, we demonstrate the utility of our on-chip engineered lymphatics to study lymphatic-immune interactions, specifically during inflammatory responses corresponding to an increased number of immune cells recruited to the lymphatics, guided by chemical gradients of lymphatic-secreted chemokines.

## 2. Results and Discussion

### 2.1. Engineered Physiological Lymphatic Microvasculature via Biochemical or Biomechanical Stimulus

To engineer physiologically-functional, human lymphatic microvasculature, we implemented a microfluidic platform that facilitated the precise spatial and dynamic control over the biochemical and biophysical factors native to the development of the lymphatics, and their physiological microenvironment. As such, the microfluidic device (**Figure 1a, b**) facilitated the compartmentalization of culture media, extracellular matrix (ECM) and cells within a 3D environment. Additionally, the PDMS-based device allowed for high-resolution imaging of microscale, cellular events via confocal microscopy. To first characterize diffusive transport, we measured the fluorescence intensity profile across the gel channel of a 70 kDa FITC-dextran as it diffused through the ECM (**Figure 1c**). The resulting profile was fitted to the 1D unsteady solution of Fick’s Second Law from which we obtained the effective diffusion coefficient (*D*) to be approximately 45 μm^2^/s, within the range of values measured *ex vivo* for the same fluorescent tracer in tissues [24,25]. Furthermore, we can use this to estimate of the timescale required for diffusion across the gel region as ~*w^2^/D*, where *w* is the width of the gel channel. Thus, a 70 kDa dextran, would diffuse across the gel in ~ 2 hrs, a small fraction of the several days required for lymphatic vascularization. This time is sufficiently short to ensure the adequate delivery of growth factors.

**Figure 1.**
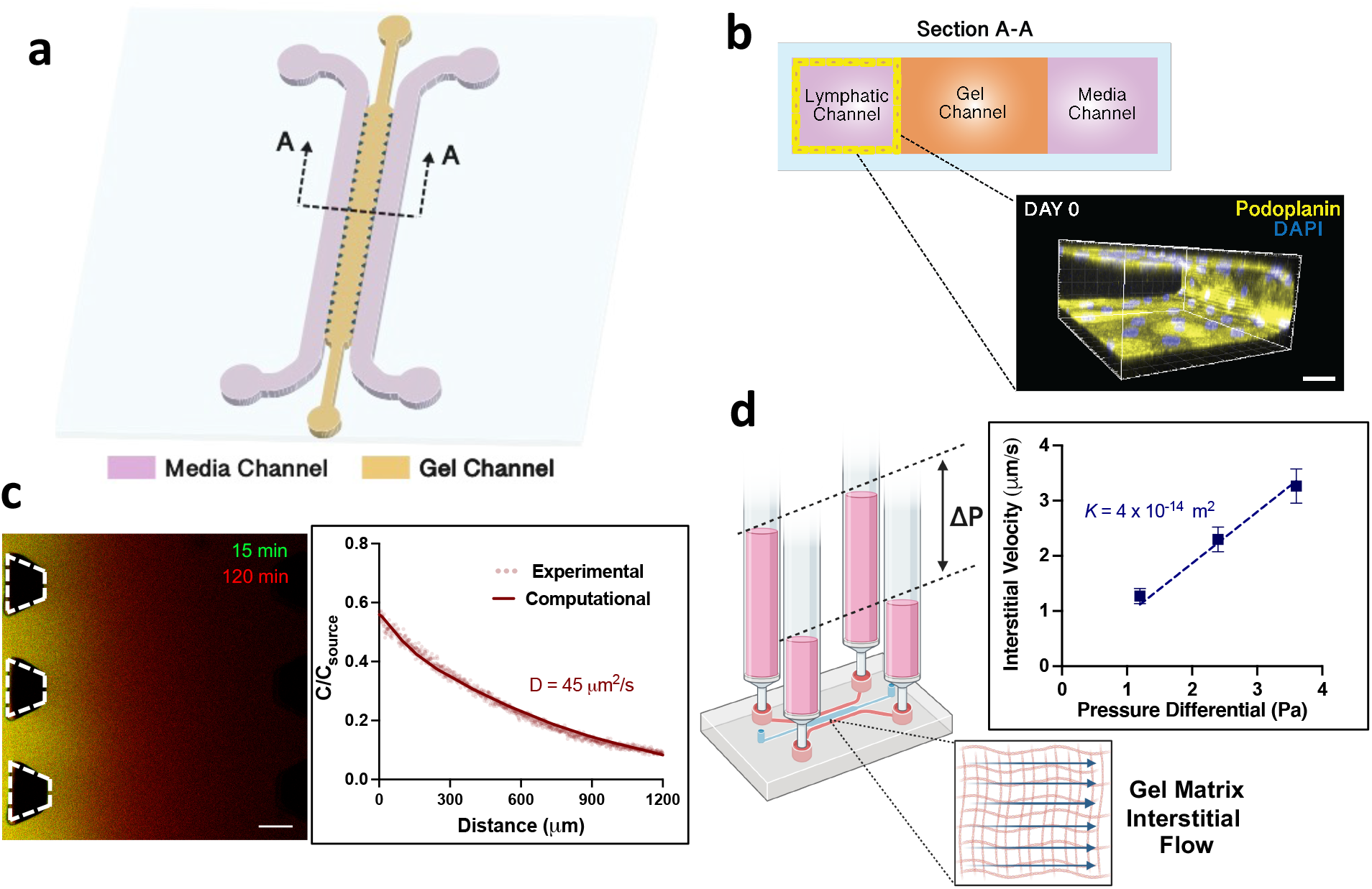
Microfluidic system for compartmentalized cell culture and precise delivery of biomolecules and fluid transport. (a) Schematic of the microfluidic device with the corresponding channels labeled. (b) Cross-sectional view of the microfluidic device. Confocal projection of a lymphatic monolayer seeded at the media channel with podoplanin staining the cell membrane and DAPI staining the nuclei. Scale bar is 100 μm. (c) Representative image of the diffusion of fluorescently-balled dextran in the microfluidic device for time intervals of 15 min (green channel) and 120 min (red channel). Scale bar is 100 μm. Intensity-concentration profile for the experimental and computational data at 120 min from which we estimated the coefficient of diffusion (*D*). (d) Schematic representation of the interstitial flow set up, where a hydrostatic pressure difference (*Δp*) drives interstitial flow across the gel matrix compartment. Interstitial flow velocity as a function of the pressure differential. The linear fit applied is based on Darcy’s law to extract the hydraulic permeability (*K*) of the gel.

In addition to the diffusive transport of growth factors, vascular fluid flow ubiquitously drives the mass transfer of biomolecules and proteins throughout tissues [26]. Pertaining to the lymphatics, the convective transport of fluid through tissue ECM (interstitial flow) and into the lymphatics facilitates the percolation and drainage of extracellular fluid and solutes to maintain hydrostatic and oncotic pressure homeostasis in tissues [27]. We are able to mimic this convective transport by introducing a hydrostatic pressure imbalance between media channels (**Figure 1d**), which drives flow across the compartmentalized ECM region. Furthermore, we measured the interstitial fluid velocity as a function of the hydrostatic pressure difference and thereby calculated the hydraulic permeability of the fibrin-based ECM (**Figure 1d**). This enables us to determine the pressure differences needed to drive interstitial fluid flow at physiological, homeostatic conditions (0.1 – 1 μm/s) [28] or, at higher velocities (~ 4 μm/s) comparable to an inflamed tissue and tumor microenvironments [29,30].

We next screened for the optimal cell culture conditions to promote the growth of 3D lymphatic capillaries resembling their native, *in vivo* morphology (**Figure 2a**). Starting from a confluent monolayer of lymphatic endothelial cells at the media-gel interface, growth factors previously identified as key mediators of developmental lymphangiogenesis (VEGF-C [31], ANG-1 [32], HGF [33]) were introduced into the opposite media channel. This generated a localized source of factors that would steadily diffuse towards the lymphatic monolayer, thus resulting in lymphatic sprouting into the central ECM compartment. The formation of lymphatic vessels was imaged for 6 days (**Figure S1**), during which morphological properties such as vessel diameter and lymphatic area of coverage were quantified. From this characterization, we sought to identify conditions that resulted in vascular morphology that overlapped with *in vivo* measured values, such as lymphatic vessel diameters within 10 - 60 μm [34], and lymphatic projected area coverage of 15 – 30 % [35–39]. The addition of lymphangiogenic growth factors, individually or in combination, consistently resulted in lymphatic sprouts with diameters well-within *in vivo* values (**Figure S3a**). However, we solely identified the simultaneous addition of all the growth factors as the optimal condition to attain the adequate range of physiological lymphatic vascular coverage within in our system (**Figure 2bi, c**). We also performed additional experiments with different combinations of the growth factors (**Figure S4**), from which we again validated that the implementation of all three growth factors is the optimal condition.

**Figure 2.**
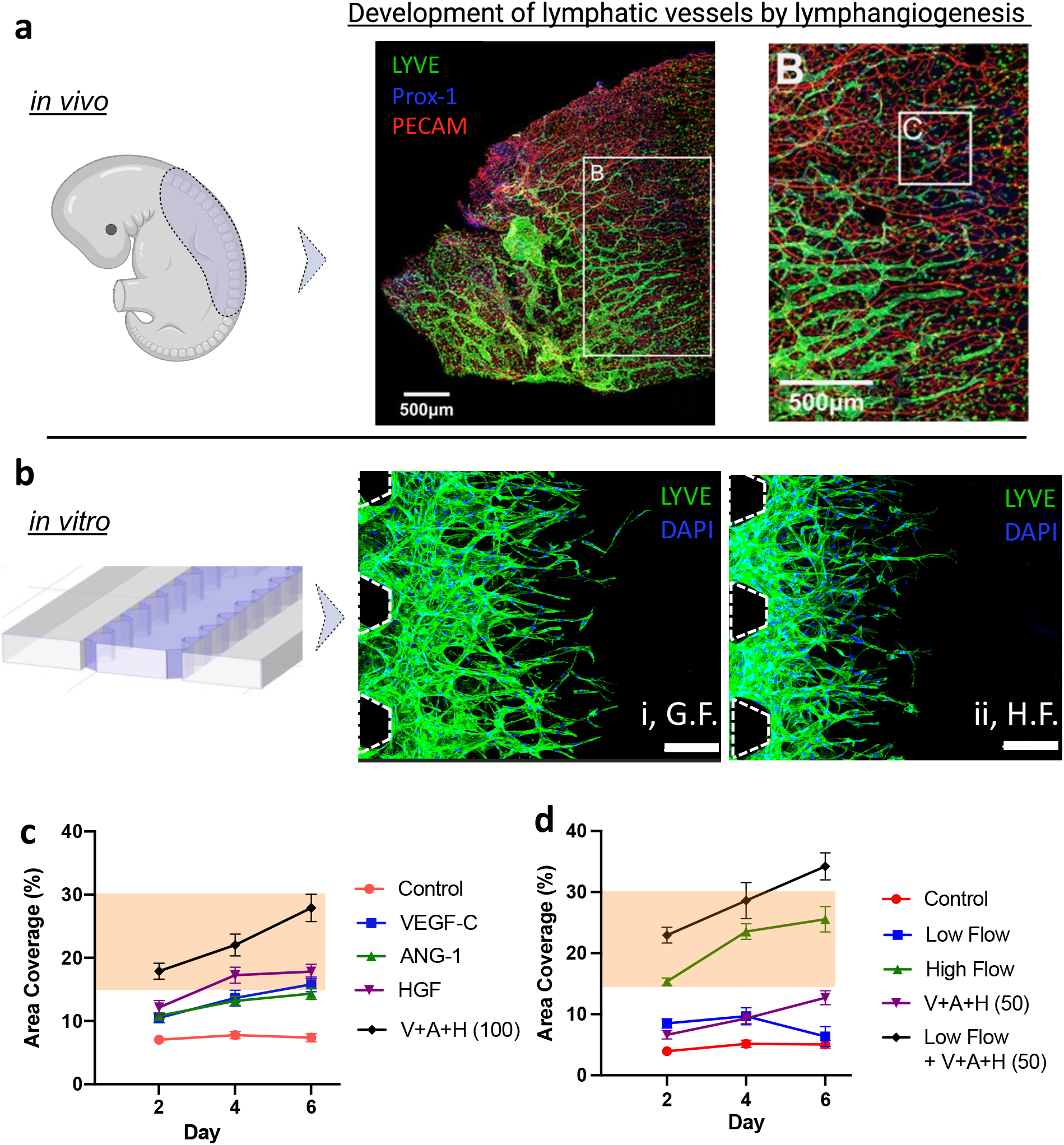
Tissue-level reconstitution of lymphangiogenesis via on-chip biochemical or biomechanical stimulus. (a) *In vivo* model of embryonic lymphangiogenesis in the dorsal skin section of a mouse embryo. Confocal images correspond to a whole-mount anterior dorsal skin (blue region in schematic diagram) with PROX-1 (blue) and LYVE1 (green) lymphatic specific markers, and blood vascular marker PECAM1 (red). Reproduced with permission via CCC [88]. (b) Microfluidic-based *in vitro* model of lymphangiogenesis, blue region in the schematic diagram highlights the ECM compartment where lymphatic vasculature is grown via growth factors (G.F.): VEGF-C, ANG-1 and HGF (i) or pathological levels of interstitial fluid flow (ii, H.F.). Confocal images depict LYVE and DAPI staining as green and blue, respectively. Scale bar is 200 μm. Quantitative analysis of lymphatic microvascular area of coverage under biochemical stimulus with growth factors (c) and biomechanical stimulus by interstitial flow at different velocities (d). Highlighted regions correspond to *in vivo* values.

In an alternate approach, we exploited interstitial flow to stimulate the growth of lymphatic capillaries based on previous work showing that sprouting is induced against the direction of interstitial flow as it passes from the extracellular space towards the vascular compartment[14,40]. We considered two interstitial flow velocities: a low flow regime corresponding to homeostatic, physiological conditions (0.1 – 1 μm/s) and a high flow regime of pathological nature (>3 μm/s). Both the low and high interstitial flow velocities elicited lymphatic sprouting during the initial days of culture, however, the extent of vascular invasion was significantly greater under high flow-conditions (**Figure 2d, Figure S2**). Higher interstitial flow velocities better mimicked the desired area of coverage for *in vivo* restitution than slower flow regimes. Thus, biomechanical stimulus, imparted by pathological-levels of interstitial flow, can be exploited as a means to generate tissue-scale lymphatic vasculature (**Figure 2bii, d**). This finding is in line with *in vivo* pathological microenvironments (i.e., a developing tumor or an inflamed wound site) where a buildup of interstitial fluid pressure, from a leaky blood vascular endothelium, leads to higher interstitial fluid flow towards the lymphatics, thus evoking lymphangiogenesis [41]. We also considered the synergistic effects of biochemical and biomechanical stimuli using a lower dose (50 ng/mL) of lymphangiogenic growth factors coupled with low interstitial flow (0.1 to 1 μm/s). Interestingly, this approach led to excessive lymphangiogenic activity resulting in hyper-physiological values for the area coverage by the grown lymphatics (**Figure 2d**). Moreover, all the diameters remained within range of *in vivo* values (**Figure S3b**), further validating this approach to grow lymphatic capillaries of appropriate length scale. We thus identified high interstitial flow by itself as best able to achieve physiologically-relevant lymphatic microvasculature and used this in all subsequent experiments.

From the acquired confocal images, we are able to affirm blind/blunt-ended, small-scale (~20 μm), 3D and lumenized structures that are more representative of their *in vivo* counterpart (**Figure 2a, b; S5a**). Furthermore, the engineered vasculature also expressed lymphatic-specific markers (**Figure S5b**) such as lymphatic vascular endothelial receptor-1 (LYVE-1) [42], and upregulated transcriptional factor PROX-1 [43].

### 2.2. Engineered Lymphatics Exhibit Solute Drainage Rates Comparable to *in vivo*

The lymphatic system serves an integral role in maintaining tissue homeostasis by clearing excess fluid, plasma proteins, pathogenic agents (bacteria and antigens) and endo-/exogenous carriers (vesicles/exosomes, therapeutics) from the peripheral tissues into the systemic circulation [27] (**Figure 3a**). Typically, the absorption of these factors is initiated by their exit from the blood vasculature followed by their transport through the tissue interstitium. This movement through the interstitial space is guided mostly by convection that transports the diluted factors towards the lymphatics, due to their lower intraluminal pressure [44]. *In vivo* studies quantifying lymphatic drainage have implemented a similar methodology as the clinical imaging technique known as lymphoscintigraphy [45], where the clearance of an injected radiolabeled or fluorescent tracer from the interstitial space is monitored as it is collected by the lymphatics. Such measurement has been widely implemented for various *in vivo* studies along different dermal sites and with tracers of distinctive molecular weights and electrochemistry, thus providing us with a range of *in vivo* clearance rates, ~ 0.02 - 0.005 min^−1^ [46–51].

**Figure 3.**
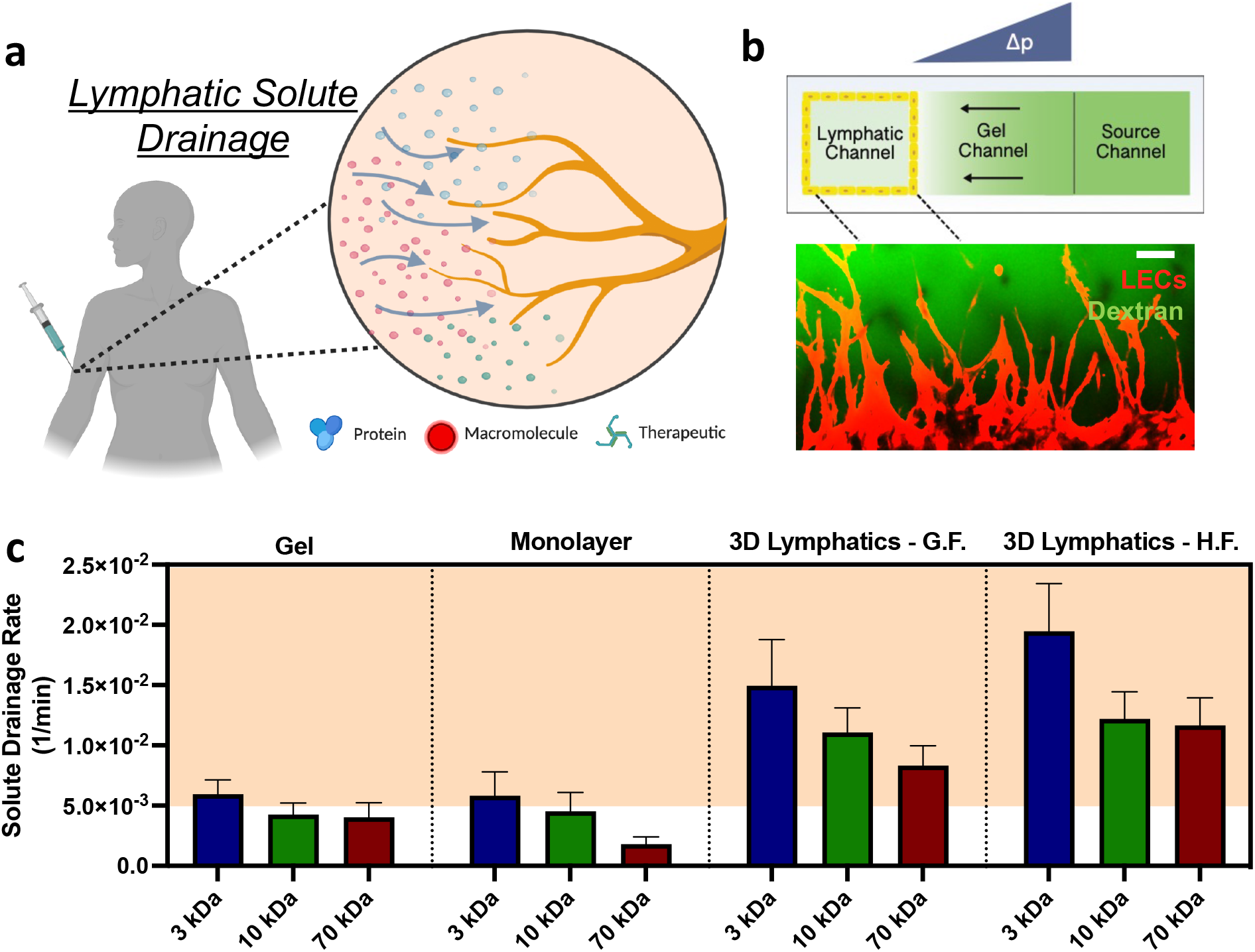
Functional characterization of physiological solute drainage by on-chip lymphatics. (a) Schematic underlying the ubiquitous physiological drainage functionality by lymphatic *in vivo*. (b) Schematic *in vitro*-based solute drainage assay within our microfluidic system to measure the drainage rate via fluorescence signal at the lymphatic media channel. Representative confocal image of the drainage assay where the lymphatics are depicted by their RFP expression (red) and the fluorescent dextran tracer (green). Scale bar is 100 μm. (c) Quantification results of solute drainage rates accordingly to the experimental condition and dextrans of varying molecular weight. From left to right, conditions correspond to: bare gel (devoid of cells), lymphatic monolayer, 3D lymphatics grown with growth factors (GF), and 3D lymphatics grown with high-levels of interstitial flow (HF). Highlighted regions correspond to *in vivo* values.

To measure lymphatic clearance in our *in vitro* engineered lymphatics, we implemented a similar approach (**Figure 3b**), previously reported by Tien and colleagues [20]. A set of four experimental conditions were studied which included 3D engineered lymphatics via optimized protocols from our previous section, implementing either growth factors or high interstitial flow. In addition, two control conditions were included corresponding to an acellular gel system, and samples with a lymphatic monolayer cultured under no angiogenic stimuli, thus the lymphatic endothelial cells mostly remained at the gel-media interface.

From the overall experimental measurements for solute drainage rates (**Figure 3c**), a series of interesting trends can be observed between systems and varying molecular weight of the tracer. For the gel system, solute drainage rates fell within or below a value of 0.005 min^−1^ which barely recapitulates *in vivo* measured rates of solute drainage. Interestingly, if we take the inverse of the average measured rate (~0.005 min^−1^) a time of 200 min is obtained which represents the timescale for the solutes to be drained through this bare gel system. This timescale is in agreement with the scaling analysis: 1200 μm / 0.1 μm/s (*w/v*) which is the convective timescale for transport across the gel width. However, given the low fluid velocity in this system, diffusive transport could also play a significant role in the migration of the solutes. To characterize the relative importance between diffusive and convective transport, we calculate the Peclet number (*Pe*), a ratio of diffusive to convective time scales:

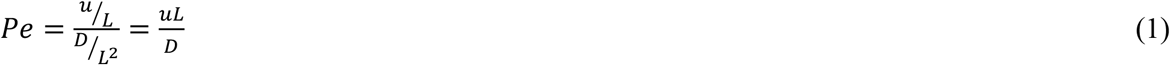

where *u* indicates the local velocity, *L* is the characteristic length of the system, and *D* denotes the diffusion coefficient of the solute. Considering the fluid velocity and *w* as the appropriate length scale, we find that *Pe* ranges from 8 – 26. Thus, transport within the gel is dominated by convection.

As we turn our attention to the lymphatic monolayer system, we observe a consistent drop in the measured solute drainage rates as we increase the molecular weight of the tracer (**Figure 3c**). This trend is similarly observed in previous experiments studying blood microvascular systems [52,53], in which both the diffusive and convective transport rates decrease with increasing tracer molecule size as a result of the inherent difficulty of larger molecules to pass between the small dimensions of the endothelial intercellular space [48]. Hence, the introduction of the lymphatics as a monolayer in our microfluidic system imposes an additional barrier to solute transport which effectively reduces the solute drainage rates to sub-physiological levels.

Evaluation of the solute drainage rates for the 3D lymphatics, engineered by angiogenic induction, with either growth factors or high interstitial flow, confirms their physiological functionality for solute drainage as we compared the measured values to *in vivo*-based measurements of solute clearance rates. Furthermore, as we increase the molecular weight of the dextran, physiological levels of interstitial solute clearance are still achieved by the engineered lymphatic microvasculature, conversely to the other systems (bare gel and lymphatic monolayer).

We also examined the drainage rates for more biologically relevant proteins, avidin and albumin. The former is extensively implemented for therapeutic particle conjugation due to its highly functional and stable affinity interaction for desired molecular targets [54], while the latter is the most ubiquitous plasma protein responsible for maintaining oncotic pressure homeostasis in the blood stream [55]. The experimental measurements further validated that the bare gel or lymphatic monolayer system failed to recapitulate physiological ranges of protein drainage rates (**Figure S7**). In fact, both proteins were cleared at similar rates as the fluorescent dextran of similar molecular weight (70 kDa). However, systems that incorporate 3D lymphatic microvasculature recurrently exhibit protein drainage rates comparable to the *in vivo-*measured values (**Figure S7**).

Beyond validating the physiological drainage functionality of our engineered lymphatics, we sought to elucidate the fundamental transport phenomena that gave rise to these distinctive drainage rates. For this we implemented both computational models and scaling arguments (Supplementary Information) from which we underline how 3D lymphatic vessel networks exhibit convective-dominant transport that ensures physiological lymphatic drainage. Such results provide theoretical insight into how we should rationally engineer 3D lymphatic vasculature to recapitulate physiological drainage.

### 2.3. Immune Recruitment by Engineered Lymphatics is Coordinated by Inflammatory-induced Chemotactic Signals

In addition to regulating the transport of diluted proteins and other macromolecules in the interstitium, the lymphatic vasculature is essential for various immune cell trafficking events during host immune responses [56]. In the event of tissue infection and inflammation, local stromal cells are activated by pathogenic signals, such as bacterial lipopolysaccharide, and respond with the release of inflammatory cytokines including tumor necrosis factor alpha (TNF-α), transforming growth factor-β (TGF-β), interleukins, amongst others [57]. Once the tissue site is primed by pro-inflammatory signals, innate and adaptive immune cells are recruited by blood and lymphatic endothelial-secreted chemokines which then initiates immune activation and response [23,57] (**Figure 4a**). Thus, in order to adequately study the signaling and migratory events that lead to the recruitment of immune cells to the lymphatics, the full range of fluid, protein and cellular transport phenomena need to be recapitulated in *in vitro* systems.

**Figure 4.**
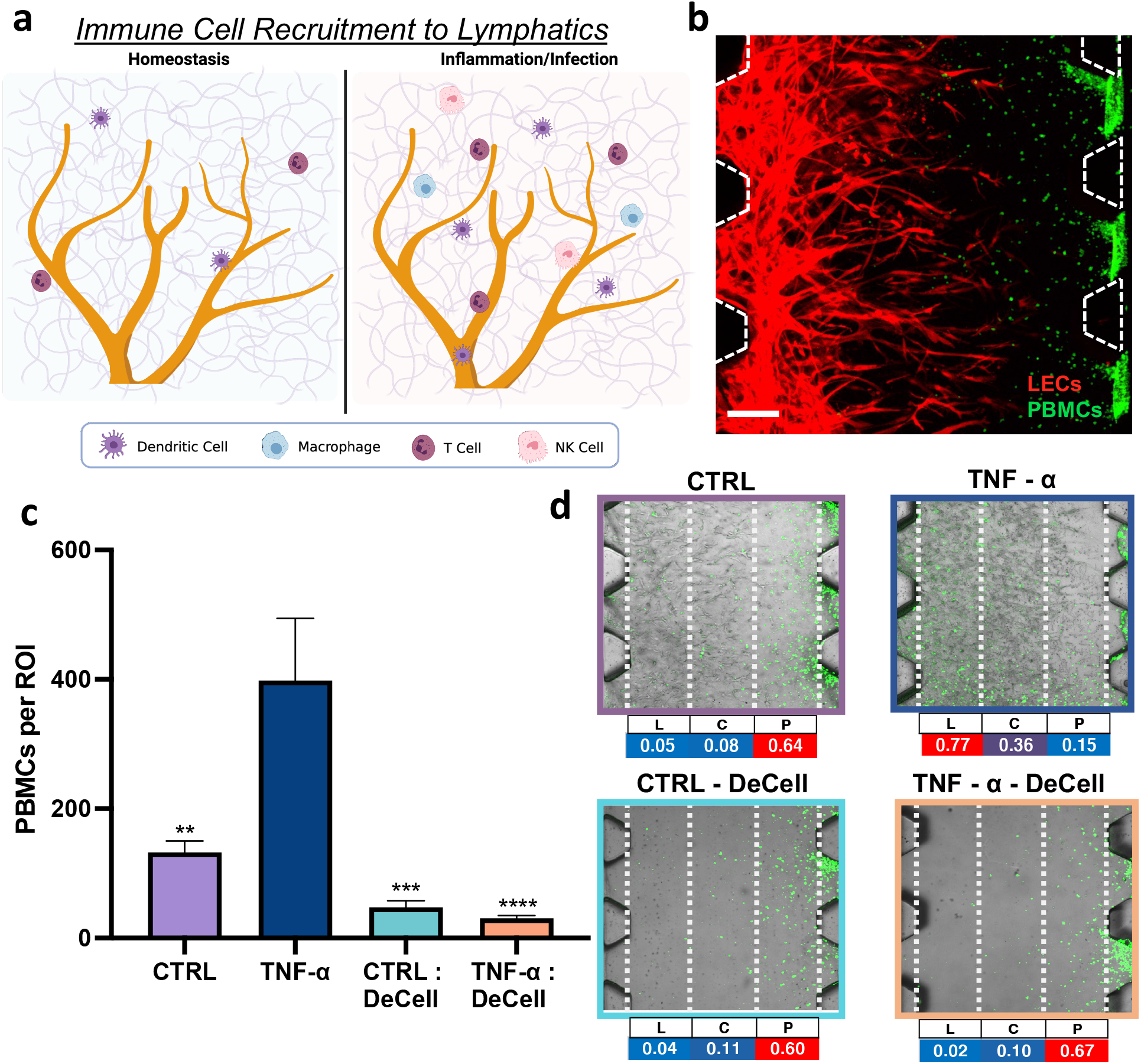
Recapitulating pathological immune cell recruitment by engineered lymphatics. (a) Schematic depiction of immune surveillance and recruitment in the lymphatic microenvironment accordingly to the tissue state. (b) PBMCs infiltration assay where the PBMCs (green) are introduced into the adjacent media channel under a hydraulic pressure difference inducing pathological flow towards the lymphatics (red). Scale bar is 200 μm. (c) Quantitative analysis from the PBMCs infiltration assay performed on high-flow engineered lymphatics with certain devices stimulated with TNF-α and/or decellularized. Statistical significance is reported with respect to the TNF-α stimulated lymphatics. (d) Representative images and quantitative heat map analysis corresponding to PBMC distribution (green) within the gel region for each experimental condition. The table columns for L, C and P correspond to the gel region with lymphatics, central gel region and region where the PBMCs are introduced, respectively.

In addition to modeling lymphatic drainage of interstitial proteins, our platform is amenable to recreating the pathological microenvironment that facilitates the intravascular recruitment of immune cells by the lymphatics. As such, immune cells can be introduced into the system at the adjacent media channel while establishing the appropriate hydraulic pressure difference across the gel compartment to impart pathological interstitial flow towards the lymphatic channel (**Figure 4b**). Additionally, recombinant-human cytokines can be introduced into the system to mimic stromal secreted inflammatory factors. As a model for immune cell response, we isolated human peripheral blood mononuclear cells (PBMCs) which includes a broad population of immune cells that are naturally found in the systemic circulation, and are known to extravasate and migrate into the lymphatic periphery. In initial experiments, PBMCs consistently migrated into the opposite channel at higher numbers for devices incorporating lymphatics (high-flow stimulated, growth factor-grown and monolayer), in comparison to the bare gel system (**Figure S14**). Given such similar trends for lymphatic-based systems, we focused the rest of our studies on solely high-flow engineered lymphatics.

To validate the increased lymphatic recruitment of immune cells under inflammatory conditions, devices with high flow-engineered lymphatics were pre-treated with TNF-α prior to PBMC perfusion, in parallel to a set of devices without TNF-α exposure which served as the control group for this experiment. Additionally, based on our solute drainage measurements, we also considered decellularized samples to examine if the increased infiltration is solely due to changes in the physical structure of the matrix, by the invading lymphatic sprouts, that could then facilitate the migration of the immune cells. Results from this set of experiments validated that, under inflammatory stimulus, lymphatics significantly increase the recruitment of PBMCs (**Figure 4c**). Additionally, decellularized samples consistently exhibited lower infiltration numbers regardless if the devices were preconditioned, or not, with TNF-α. To further examine the preferential migration and infiltration of PBMCs, confocal imaging of the gel region was done to visualize the spatial distribution of PBMCs immediately after performing the infiltration assay. Both images and quantitative data indicate that PBMCs where preferentially localized closer to the media region where they are introduced, for all conditions except the TNF-α-stimulated lymphatics (**Figure 4d**). For the latter, the highest population of PBMCs corresponded to the region co-localized with lymphatic vessels. Such data suggest that, under inflammatory stimulus, our on-chip lymphatics provide directional cues that elicit the preferential migration and infiltration of PBMCs.

Efforts to understand the basis of immune cell migration to specific sites during immune response, have elucidated the role of chemokines as homing molecules for immune recruitment [58]. These chemotactic cytokines guide immune cells to different sites and at different steps during immunogenic response which requires the careful coordination and delivery of cell-secreted ligands to the specific immune cell surface receptors [59] (**Figure 5a**). Of the most studied immune chemotactic pathways during inflammation are: lymphatic secreted ligands CCL21 and CCL19 that attract immune cells via their CCR7 receptor during various pathological events including immunogenic response and cancer metastasis [60–62], and stromal/endothelial secreted CXCL12 that coordinates the homing of immune cells for immune maintenance and development by their CXCR4 surface receptor [63,64]. To determine if these corresponding receptors are elicited during inflammatory immune recruitment in our tissue engineered system, we performed receptor blocking experiments by incubating PBMCs in a buffer solution (containing CCR7 and/or CXCR4 antibodies, or control isotypes), prior to performing the infiltration assay with high-flow grown lymphatics stimulated with TNF-α. For this panel of experiments, we observed a consistent and significant decrease in the number of infiltrated PBMCs when their cell surface receptors, CCR7 and/or CXCR4, are functionally blocked, compared to the IgG isotype control, despite being introduced into devices with inflammatory-stimulated lymphatics (**Figure 5b**). Furthermore, analysis of the spatial distribution of PBMCs within the gel region reveals that solely for the IgG isotype control samples, the PBMCs preferentially migrated and co-localized at the inflamed lymphatics region (**Figure S15**). All other samples with neutralizing antibody treatment displayed higher PBMC numbers closer to the media channel. Thus, the CCR7 and CXCR4 immune cell surface receptors, responsible for immune homing, are elicited in our engineered pathological model for immune cell recruitment by the lymphatics in response to local inflammatory stimulus. However, we anticipate that additional immune cell receptors are co-activated during the recruitment of PBMCs during infiltration. As such, future studies to interrogate activated immune chemotactic receptors during lymphatic recruitment can be performed with our platform.

**Figure 5.**
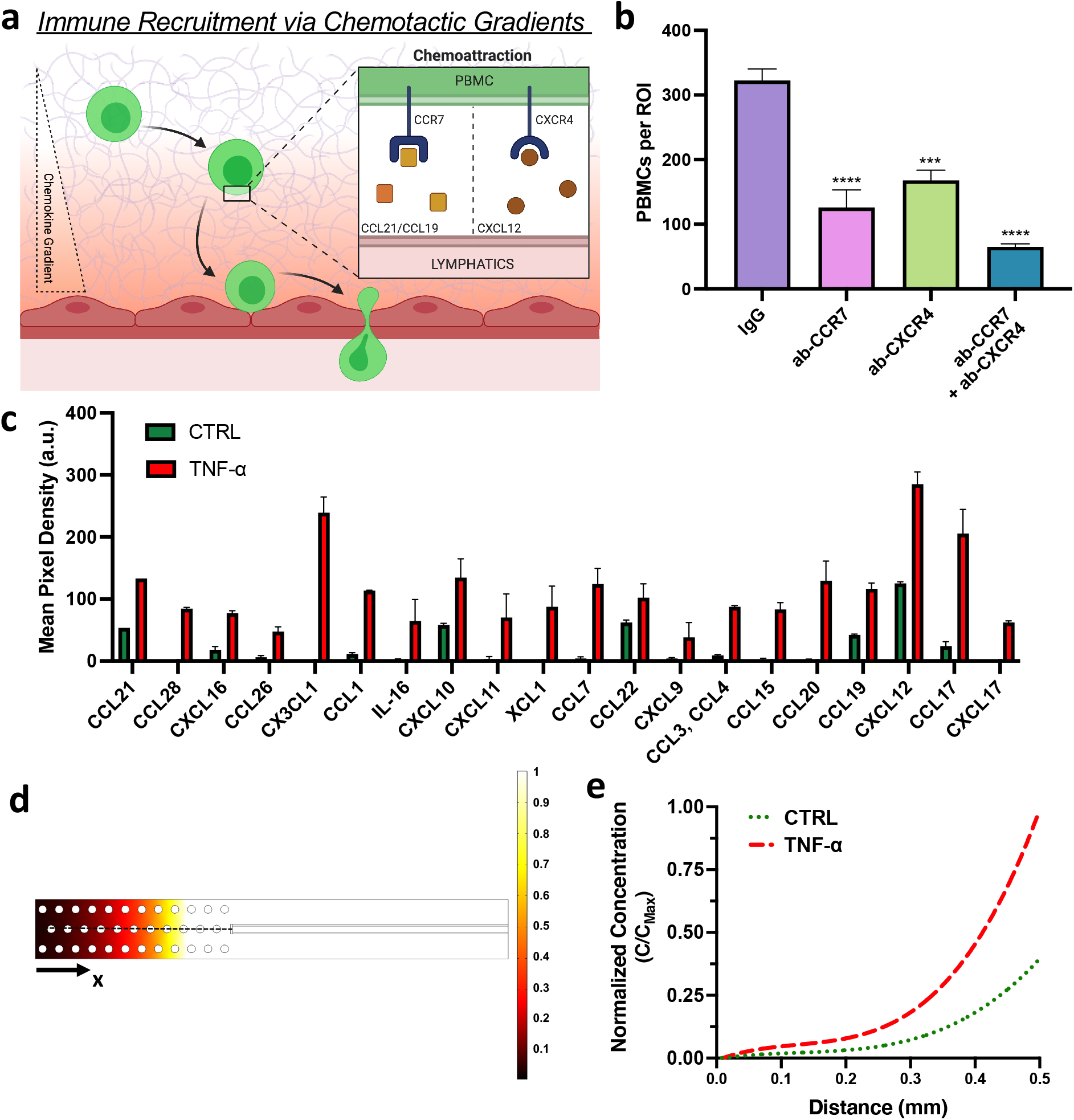
Flow-induced concentration gradient of chemotactic factors facilitate immune recruitment by on-chip lymphatics. (a) Schematic diagram on the chemotactic signaling axes that recruit immune cells via concentration gradients of the corresponding chemokines during an inflammatory response. (b) Quantitative analysis from the PBMCs infiltration assay performed on high-flow engineered lymphatics stimulated with TNF-α and with corresponding conditions of PBMCs reconstituted in neutralizing antibodies or IgG isotypes. Statistical significance is reported with respect to the experimental condition with PBMCs reconstituted in IgG isotypes. (c) Quantitative analysis of cytokine secretion by lymphatics grown under interstitial flow with one experimental group pre-conditioned with TNF-α prior to collecting their respective supernatant. (d) Computational model results for lymphatic chemokine transport, developed in COMSOL Multiphysics, illustrating the normalized concentration of the chemokines. Dashed line indicates the line probe from which the concentration plot (e) is generated considering differences in secretion rate by untreated (control) lymphatics and TNF-α-stimulated samples.

To comprehensively validate the increase activation and recruitment of PBMCs by lymphatic-secreted chemokines during inflammation, we quantified cytokine expression by collecting the supernatant from lymphatic cells cultured under the similar experimental conditions. Quantitative analysis of the cytokine intensity between both experimental conditions revealed a consistent increase in chemokine secretion by the TNF-α-conditioned lymphatics, with an approximate 2.5-fold increase for CCL19, CCL21 and CXCL12 (**Figure 5c**). Conclusively, the immune surface receptors CCR7 and CXCR4 mediate the increase infiltration of PBMCs, while lymphatics secrete corresponding ligands that activate these chemotactic receptors within our inflammatory immune recruitment model.

When considering the physio-/pathological environment by which these chemokines are secreted into the extracellular space, most of them remain soluble or, alternatively, bind to the matrix, which would result in concentration gradients [65–67]. Thus, facilitating the directed migration and recruitment of the immune cells to the specific tissue or vasculature via graded chemical concentrations [68,69]. In order to emulate the paracrine signaling that leads to the recruitment of immune cells in our engineered system, we sought to demonstrate that these chemotactic conditions could be established and maintained even during interstitial flow toward the lymphatics. Given the lack of experimental techniques to precisely visualize and quantify the spatial distribution of chemotactic factors in real-time, we extended upon our computation lymphatic drainage model, and incorporated an additional domain consisting of solid (impermeable) spheres embedded in the gel region which represented the migrating PBMCs. Based on a quasi-steady state approximation (supported by scaling arguments, Supplementary Information), we obtain a generalized picture on the distribution of chemokines which affirmed the ability of our platform to recapitulate concentration gradients of lymphatic-secreted chemokines (**Figure 5d**). In fact, numerical results indicate that the spatial distribution of the chemokine follows a concentration gradient leading towards the lymphatics (**Figure 5e**). This gradient is facilitated by competing transport where interstitial flow skews the symmetric diffusion of chemokines, thus resulting in a trail of lower concentration at farther distances from the chemokine source (lymphatics). We can provide additional quantitative basis to this phenomenon by considering the scaling analysis provided by the Peclet number, as described in previous sections. The corresponding value for the Peclet number is approximately 10, which is within the characterized range of flow-induced transcellular gradients of cell secreted factors [70]. However, an additional aspect is to be evaluated which is the relative difference between homeostatic and inflamed lymphatics with varying rates of chemokine secretion. Despite not having experimental characterization of chemokine secretion rates, we do have quantitative insight on the relative increase of secreted chemokines between the TNF-α-stimulated lymphatics and the unstimulated condition. From the cytokine array analysis, we found that the chemokines of interest are secreted at a 2.5x increased rate for inflammatory-stimulated lymphatics. We applied this relative increase to the flux of species by the lymphatic sprout in our model to which we found a significantly steeper profile in the concentration gradient of the chemokines (**Figure 5e**). As extensively studied *in vitro* [71–73], such increase in spatial gradients of chemotactic factors results in faster and more persistent migration patterns by immune cells, which is in line with our experimental findings that higher numbers of immune cells infiltrate our inflamed lymphatics model.

## 3. Conclusion

In this work, we address the need for a physiologically relevant *in vitro* model that recapitulates critical aspects of the *in vivo* lymphatics, including their tissue architecture and physio-/pathological transport functionality. For this purpose, we implemented microfluidic-based cell culture system that allowed us to compartmentally culture lymphatic endothelial cells under the appropriate biological stimuli to induce their self-organization into *in vivo*-like capillaries. Furthermore, we leveraged the capabilities of our microfluidic system to recreate the biological transport of fluid, interstitial solutes and immune cells which allowed us to perform functional assays with the on-chip engineered lymphatics. On the basis these studies, we successfully recapitulated key aspects in the physiological drainage functionality of lymphatics and the immune-lymphatic pathological response. These results further support the use of our microphysiological platform for pre-clinical applications that would be technically challenging to perform with *in vivo* models, such as characterizing lymphatic absorption of subcutaneously-delivered therapeutic antibodies and screening immunogenic response of genetically-modified immune cells that transit across the lymphatic interface to reach their corresponding cellular target.

## 4. Experimental Methods

### Microfluidic Device Design and Preparation

The microfluidic device was based on earlier designs from our lab [74–76] with three, parallel fluid channels. The middle channel is lined by a series of trapezoidal posts at the edges adjacent to the other fluid channels. These provide adequate surface tension to facilitate the compartmentalization of the injected extracellular matrix in the middle region. The side channels are then utilized as medium channels that allow the exchange of nutrients and metabolic waste, as well as to supply growth factors at specific boundaries of the gel region. Additionally, the length scales of the system were optimized to facilitate lymphatic vascularization in the middle, gel channel. The distance between media channels, which defines the width of the gel region, is 1.2 mm -- sufficiently short to allow the steady diffusion of growth factors from one media channel to the other within several hours, while providing sufficient distance for the lymphatic cells to generate vascular sprouts in the gel region [74]. On a similar basis, a width of 1 mm was set for the media channels. The device height, 300 μm, was chosen to maximize the 3D space of lymphatic vascularization, while still facilitating high-resolution confocal imaging. Finally, the length of the media-gel interface was extended to 1cm to approach a tissue-relevant scale while still maintaining a small device footprint.

Microfluidic devices were fabricated by soft lithography from SU-8 coated silicon molds similarly to previous protocols [77]. Briefly, molds were prepared by photopolymerizing a 300 μm thick SU-8 photoresist (Micro-Chem, USA) on the silicon wafer. After developing the SU-8 layer, the wafer was silanized overnight in a vacuum desiccator to facilitate the passivation of the surfaces, thus preventing PDMS adhesion during removal. A 10:1 mix of PDMS (Sylgard 184, Ellsworth Adhesives, USA) and curing agent was then poured onto the mold, allowed to degas in a desiccator for ~30 min, and polymerized at 70 °C for at least 2 hrs. PDMS was then removed from the mold and cut into individual devices. Scotch tape was used to further clean the surface of the device removing dust and particulates. To allow upper access to the fluid channels, ports were punched using a 1.2 mm biopsy punch for the gel channel, and a 6 mm or 4 mm biopsy punch for the media channels in devices used to grow the lymphatic microvasculature via growth factors or interstitial flow, respectively. After dry sterilization of the devices, the surface was treated with plasma (Harrick Plasma, USA) for 90 seconds, and then bonded to a coverslip slide. After plasma bonding, devices were left overnight to recover hydrophobicity and kept sterile until use.

### Cell Culture

Human dermal lymphatic microvascular endothelial cells (CC-2543, HDLMEC, Lonza, USA) were cultured in Vasculife Endothelial Medium (Lifeline, LL-0003) supplemented with 6% FBS (Invitrogen). HDLMEC were transduced to express cytoplasmic RFP using LentiBrite™ RFP Control Lentiviral Biosensor (EMD Millipore, 17-10409) as described by the vendor. Cells were cultured at 37 °C and 5% CO_2_ in a humidified incubator with media replacement every second day. HDLMEC were used in experiments before reaching confluence, between passages 6-8.

### Growth Factors, Fluorescent Tracers and Antibodies

All reagents were reconstituted to a stock solution as recommended by the corresponding vendor, and then diluted to desired concentrations. Vascular endothelial growth factor-c (VEGF-C, R&D Systems), angiopoietin-1 (ANG-1, R&D Systems) and hepatocyte growth factor (HGF, Peprotech) were all used at a 100 ng/mL dilution in the cell culture medium for specified experimental conditions. Immunostaining of the lymphatic vasculature was performed using PE-conjugated anti-human podoplanin (Biolegend), Alexa Fluor® 647-conjugated anti-human lymphatic vessel endothelial receptor-1 (LYVE-1 ,R&D Systems) or Alexa Fluor™ 488-conjugated anti-human laminin (R&D Systems) to image the cell surface and DAPI (Invitrogen) or Alexa Fluor™ 594-conjugated anti-human Prox1 (Biolegend) to image cell nuclei at a 1:100 and 1:1000 dilution from stock in washing buffer (0.5% BSA/DPBS), respectively. For diffusive and convective transport measurements, fluorescent dextran: 3 kDa-Cascade Blue™ (Thermofisher), 10 kDa-Cascade Blue™ (Thermofisher), or 70 kDa-Fluorescein (FITC, Thermofisher) was supplemented in the cell culture medium at concentration of 100 μg/mL. In a similar set of experiments, Fluorescein-conjugated avidin (Thermofisher) or Alexa Fluor™ 647-conjugated albumin from bovine serum (Thermofisher) was introduced into the system at a concentration of 100 μg/mL. For neutralizing antibody experiments, PBMCs were incubated in a buffer solution with their respective antibody blocker at a concentration of 5 μg/mL: human CCR7 antibody (MAB197-SP, R&D), human CXCR4 antibody (MAB172-SP, R&D), and a combine human IgG_2A_ and IgG_2B_ isotype control (MAB003 and MAB004, R&D).

### Device Seeding and Microvascular Culture

Fibrinogen from bovine plasma (Sigma) was dissolved for at least 3 hrs at 37 °C in Dulbecco’s Phosphate-Buffered Saline (DPBS, Lonza) at twice the final concentration, 5 mg/mL. A thrombin (Sigma) stock solution was made at 100 U/mL in 0.1% w/v bovine serum albumin (BSA) solution and stored at −70 °C. Thrombin was diluted in Vasculife Endothelial Basal Medium to a concentration of 4 U/mL. Solutions were mixed via pipetting, over ice, in a tissue culture hood at a 1:1 ratio to produce a fibrin solution with a final fibrinogen concentration of 2.5 mg/mL. The mixture was then pipetted into the device using the gel filling ports. Devices were placed in a humidified enclosure and allowed to polymerize at room temperature for 15 min.

Human Plasma Fibronectin (EMD Millipore) was diluted to a concentration of 100 μg/mL in DPBS, prior to being injected into one of the media channels where the HDLMEC would be seeded in order to facilitate their adhesion to the walls of the device. While the devices were left incubating with the fibronectin solution for at least 30 min, HDLMEC were trypsinized (Lonza, USA) and resuspended to a concentration of 3 × 10^6^ cells/mL. After incubation, fresh media was introduced into the fibronectin coated channels and aspirated, followed by perfusion of 30 μL of the cell suspension into the channel. Immediately after cell seeding, devices were tilted by approximately 120° and incubated for 15 min to facilitate the adhesion of cells on the gel-media interface. Subsequently, devices were returned to their original position and fresh media was supplemented into the remaining media channel devoid of cells. A pressure difference of ~10 Pa was established between media channels with flow directed from the lymphatic media channel towards the opposite media channel to further assist the accumulation of cells at the interface. After 24 hrs of culture under these conditions, a confluent monolayer of lymphatic endothelial cells forms at the gel media interface and the remaining unattached cells are aspirated.

Following the formation of a confluent lymphatic monolayer (**Figure 1b**), ~350 μL plain Vasculife Endothelial Medium was replenished in the lymphatic media channel. To stimulate lymphatic sprout formation into the 3D gel region, ~350 μL of cell culture medium supplemented with the specified growth factor was added to the adjacent media channel, and replenished on a daily basis for up to 6 days. Previous dose-response experiments were also performed (data not shown) for the concentration of growth factors, from which we identify a concentration threshold (100 ng/mL), above which lymphatic sprouting is not significantly improved during the course of the experiments. For experiments where interstitial flow was implemented to stimulate lymphatic sprouting, a pressure head difference was applied using disposable 10 mL luer-slip syringes (Kinesis) cut at the 4 mL label, then dry sterilized. Once the lymphatic monolayer was generated, syringes were gently press-fit into the media ports and media was added to the reservoirs accordingly to the desired pressure difference (either 100 or 10 Pa for high and low flow, respectively) and interstitial flow velocity (**Figure 1d**). The hydraulic pressure head difference was reestablished on a daily basis up to 6 days, with minimal pressure loss over a 24-hr period (~10%).

### Immunofluorescence Staining, Imaging and Quantification

Growth of the lymphatic vasculature was measured every other day over a course of 6 days by taking epifluorescence images of their RFP signal on a Nikon Eclipse Ti-S (Nikon Instruments, USA) at 4x with a numerical aperture of 0.13. These parameters permit an imaging thickness of ~25 μm [78], which is similar to the thickness of histological sections used to quantify *in vivo* lymphatic morphology [79]. The Lymphatic Vessel Analysis Protocol-plugin in ImageJ (NIH) was utilized to measure morphological properties under the same protocol as implemented for lymphatic capillaries from tissue cryosections [80]. The area of coverage was quantified from binarized images that showed the relative area within the gel invaded by the lymphatics. Vessel diameter was measured at 2 to 3 locations along each sprout, for a total of 5-20 sprouts per imaged area (depending on the number of available sprouts to measure accordingly to experimental condition). For immunofluorescence imaging, cells were fixed with 4% paraformaldehyde (Electron Microscopy Sciences) for 15 min, followed by permeabilization with 0.01% Triton™ X‐100 (Sigma) for 10 min. Subsequently, blocking was performed with 5% BSA (Sigma) and 3% goat serum (Sigma) for 1 hr at room temperature. All the reagents were diluted in DPBS. Cells were then incubated overnight at 4°C with a corresponding protein-antibody of interest. After incubation, the samples were washed 5x with washing buffer and stored at 4°C. Confocal images were acquired with IX81 microscope (Olympus) equipped with Fluoview FV1000 Software (Olympus).

### Diffusion, interstitial flow and hydraulic permeability assessment

For characterization of the intrinsic transport properties of the gel, device gel seeding was followed as specified in the previous section but without the addition of cells and left to incubate overnight. Cell culture media supplemented with 70 kDa-FITC dextran was added to one of the media channels and the middle, gel region was imaged at specified time intervals on a confocal microscope (Olympus FV-1000) with custom enclosure for temperature and atmosphere control. Fluorescent signal obtained from the images depicted changes in the concentration profile over time due to diffusive transport of the molecules across the gel. Images were taken at the midplane with a 10x objective, then analyzed using ImageJ (NIH) to extract the fluorescence intensity profile across the width of the gel channel. Computational simulations were performed to obtain the diffusion coefficient of the tracer within the gel region (Supporting Information).

To quantify interstitial fluid velocity corresponding to the established pressure difference across the gel, a variation of the Fluorescence Recovery After Photobleaching (FRAP) technique was implemented as previously reported [81]. Devices were perfused with 70 kDa-FITC dextran supplemented media and left overnight to ensure its uniform distribution throughout the gel. Subsequently, 3 to 5 small regions of interest (ROIs) 30 μm in diameter were photobleached by setting the confocal laser power to 100% for ~5 seconds within the gel matrix. Time-lapse images were captured at 10x immediately after photobleaching every 1.5 seconds. By measuring the translation of the photobleached spot centroid using the Matlab frap_analysis plugin [82], we extracted the interstitial fluid velocity at each pressure head difference. This assumes that: (1) the tracer molecule is small compared to the pore size of the fibrin matrix [83], and (2) the flow field is uniform, which is also valid since the length scale of the gel compartment is orders of magnitude greater than the distance over which the velocity varies (*w, h* ≫ *K* ^0.5^); thus, the flow velocity is nearly uniform across the gel, falling to zero over the small distance *K* ^0.5^ at the device walls [84]. This measurement was repeated as the hydraulic head was increased to obtain an experimental trend between the pressure offset (*Δp*) and interstitial flow velocity (*v*) to confirm the linear relationship (Darcy’s Law (Equation 2)), and determine the hydraulic permeability of the gel (*K*) given by:

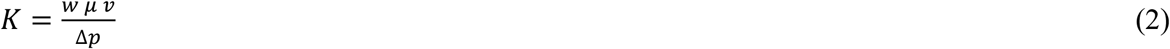

where *w* indicates the length over which the pressure drop is imposed (gel channel width), and *μ* corresponds to the fluid viscosity taken as 0.78 cP from previous studies [85].

### Lymphatic Solute Drainage Rate Assessment

A solute drainage assay was implemented based on previous *in vivo* techniques and adapted for *in vitro* models [20,86]. On day 4 post-seeding/culture, media supplemented with a fluorescent tracer was introduced into one of the media channels to establish a hydraulic pressure difference between media channels and drive flow through the gel at an average interstitial fluid velocity of ~ 0.1 μm/s, thus recapitulating physiological flows towards the lymphatics. Immediately after, a series of 4 ROIs at the opposite media channel (initially solute-free) are imaged as a confocal stack throughout the full height of the device (4 slices at 80 μm) at 10x every 2-4 min for up to 12 min to determine average fluorescent intensity over time. Additional images were acquired for the source channel, where the fluorescently-conjugated solutes are originally introduced. The average fluorescence intensity of each corresponding ROI was extracted using ImageJ-based quantification and used to calculate the solute drainage rate:

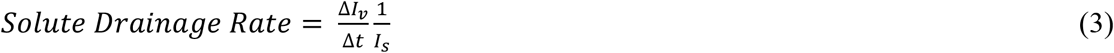

where *ΔI*_*v*_ indicates the increase in the average fluorescence intensity within the lymphatic vasculature over a time interval (*Δt*), and *I*_*s*_ is the average intensity in the source channel. A fundamental assumption imposed by this metric is the linear increase in fluorescence intensity as the tracer is drained into the lymphatic channel. Measurements from this assay by Tien and colleagues [20] and our study (**Figure S6**) validate a nearly linear trend in the fluorescence signal, corresponding to convection-dominated transport of solutes into the lymphatic channel.

For devices containing lymphatic endothelial cells, an additional solute drainage rate assessment was conducted after decellularizing the system. This was done by washing away the cells with a detergent solution of 1% Triton™ X‐100 in DPBS for 10 min immediately after the first solute drainage measurement, followed by washing the previous fluorescent tracer using DPBS for 5 min. All the washing steps were done under a slight pressure head difference (~10 Pa). Solute drainage rate measurements were repeated with decellularized devices to determine the relative difference between measured values prior and post lymphatic decellularization.

### Lymphatic Diffusive Permeability and Hydraulic Conductivity Assessment

To quantify diffusive transport across the lymphatic endothelium, a diffusive permeability assay was implemented based on previous protocols [52]. Briefly, devices were perfused with a fluorescent tracer at the lymphatic media channel, and allowed to diffuse through the endothelium while confocal z-stacks of > 100 μm thick were collected in steps of 5 μm and at time intervals of 2 min. Considering the principle of mass conservation (Equation 4), the diffusive flux (*N*_*d*_) of the solutes from the lymphatic endothelium into the gel region can be quantified by taking the temporal derivative of the total concentration (*C*) in the gel control volume (*V*_*g*_):

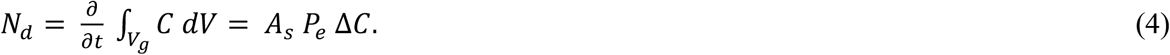

Since the total diffusive flux is driven by the concentration difference (*ΔC*) across the surface area of the vasculature (*A*_*s*_), and assuming that the fluorescence intensity is linearly proportional to the concentration of the fluorescent tracer, the diffusive permeability of the endothelium (*P*_*e*_) can be computed from [87]:

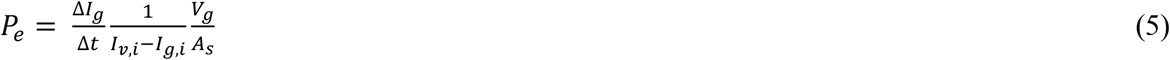

where *ΔI*_*g*_ is the increase of average fluorescence intensity in the gel region after a given time interval (*Δt*), respectively, and *I*_*v,i*_ and *I*_*g,i*_ indicate the initial, average fluorescence intensity within the lymphatic vasculature and gel volume, respectively. This measurement was repeated at 3 locations per device for fluorescent tracers of varying molecular weight.

Fluid transport across the lymphatics was characterized in terms of an endothelial hydraulic conductivity. For this assessment, interstitial flow velocities (*v*) in devices with lymphatics were measured at the same pressure head differences (*Δp*_*total*_) used for the estimation of the hydraulic permeability. Following these measurements, and recognizing that the major sources of hydraulic resistance (ratio of differential pressure and volumetric flow rate) in the system originate from the gel and lymphatic endothelium [40], the differential pressure across the endothelium (*Δp*_*ec*_) can be estimated by:

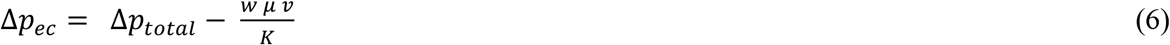

where the differential pressure across the gel is governed by Darcy’s Law, as previously described. Knowing the pressure drop across the endothelium, we can apply the Starling equation [ref] to establish the proportionality between the pressure difference and fluid flux across the lymphatics, known as the hydraulic conductivity (*L*_*p*_):

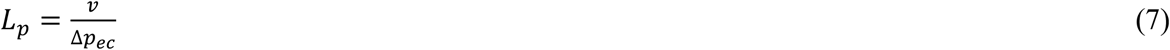

where the oncotic pressure contribution is neglected assuming nearly homogenous concentration of solutes throughout the system. This quantification was performed for each configuration of the microfluidic system (monolayer, growth factors- and interstitial flow-grown lymphatics). All transport parameters were measured and utilized for the simulations are listed in Table S1.

### Immune Cell Isolation and Recruitment Assessment

Human blood collected from healthy donors (Research Blood Components, Massachusetts, USA) with anticoagulant sodium citrate, was mixed at a ratio of 1:1 with a buffer solution, consisting of DPBS supplemented with 0.1% BSA (Sigma) and 1 mM of ethylenediaminetetraacetic acid (ThermoFisher), and carefully layered over Ficoll^®^ Paque Plus (Sigma) in a Corning 50 mL centrifuge tube. Then, the layered blood samples were centrifuged at 400 rcf for 35 min. Following centrifugation, the upper layer, consisting of plasma, was removed by aspiration, and the remaining buffy layer (containing PBMCs) was transferred to a separate centrifuge tube. Additional purification from any remaining platelets was performed by diluting the isolated PBMCs in buffer solution supplemented to the remaining volume of the 50 mL tube, and centrifuging the resuspended cells at 250 rcf for 10 min. Platelets mostly remained in the supernatant, which was aspirated, and PBMCs remained agglomerated at the bottom. This final step was repeated twice, as recommended by the standard isolation protocol to yield >90% of PBMC purity. All isolation procedures were done at room temperature. Immediately after purification, PBMCs were stained with Cell Tracker™ Green CMFDA Dye (Invitrogen) at a 10 μM concentration for 10 min, followed by washing with the buffer solution, and resuspended in cell culture media to a final concentration 1 × 10^6^ cells/mL. For receptor blocking experiments, an additional incubation step was implemented before the final resuspension step, with PBMCs incubated in the buffer solution containing CCR7 and/or CXCR4 antibodies, control IgG antibodies or no treatment for 30 min. After the final resuspension, 200 μL of the PBMC solution (~200,000 PBMCs) was perfused into the media channel devoid of lymphatic cells in devices at day 4 of cell culture. For some cases, devices were pre-treated with 20 ng/mL of TNF-α (Peprotech) overnight. Prior to PBMC perfusion, any remaining TNF-α was washed away to ensure that inflammatory stimulus was solely applied to the lymphatics. Once the PBMCs were introduced into the devices, a hydraulic pressure head was established to drive flow towards the lymphatic channel at pathological interstitial fluid velocities (~4 μm/s), as estimated in inflamed tissues [29,30]. Following 24 hrs of this assay, epifluorescence images were acquired at the lymphatic channel, and the total number of PBMCs was quantified using the TrackMate-plugin in ImageJ. Additional imaging was done with fixed devices which included confocal imaging the gel region to visualize the spatial distribution of PBMCs. Quantitative analysis of the images was performed by subdividing the gel into three equally-spaced regions along its width (Fig. 5(d)), and measuring the average fluorescence intensity of each region which correlated with the density of immune cells.

### Chemokine Secretion Analysis

Chemokine expression was quantified by collecting the supernatant from lymphatic cells cultured under experimental conditions similar to those for the immune recruitment assay. Prior to collection, lymphatic endothelial cells (~10^6^ cells) were seeded on a Corning™ Transwell Membrane Inserts with 0.4um pores (Sigma). After 24 hrs of culture a lymphatic monolayer was established on the transwell insert, and 1 mL of fibrin gel was injected on top of the monolayer and allowed to polymerized. Subsequently, cell culture media was added to the top of the gel inducing interstitial flow (~4 μm/s) towards the lymphatics with one of the transwell having cell culture media supplemented with 20 ng/mL of TNF-α. Following 24 hrs of culture under these conditions, the supernatants were collected and assayed in a Proteome Profiler Human Chemokine Array (ARY017, R&D systems) following manufacturer’s protocol. Chemiluminescent imaging was performed using Alpha Innotech (USA), and the generated images were analyzed using the Protein Array Analyzer-plugin in ImageJ.

### Statistical Analysis

Statistical analysis was done in Graphpad Prism (Graphpad, USA). All data shown represents experiments with n = 2-4 individual devices per condition. Reported values correspond to averages over these devices with error bars representing the standard error of the mean. Statistical significance was calculated with ANOVA and all tests resulting in a *p-value* less than 0.05 were considered statistically significant and were grouped as \**p* < 0.05; \*\**p* < 0.01; \*\*\**p* < 0.001; \*\*\*\**p* < 0.0001; ns (not significant), *p* > 0.05.

## Acknowledgements

The authors would like to thank Prof. Alan Grodzinsky, Prof. Ming Guo, and members from the MIT Mechanobiology lab for insightful discussions about this work. J.C.S. is supported by an NSF Graduate Research Fellowship. M.R.G. was supported by from a Canadian Institutes of Health Research Postdoctoral Fellowship. R.D.K. acknowledges funding from the National Science Foundation Science and Technology Center for Emergent Behaviors of Integrated Cellular Systems (No. CBET-0939511) and from the National Cancer Institute (No. U01 CA214381). We also acknowledge the Swanson Biotechnology Center at the David H. Koch Institute for Integrative Cancer Research for the exceptional core facilities, and the MIT Biomicro Center for providing access to COMSOL software. Schematic figures were created with BioRender.

