## Supplementary Information for "On-chip engineered human lymphatic microvasculature for physio-/pathological transport phenomena studies"

### Supplementary Text

#### Finite Element Computational Modelling

COMSOL Multiphysics (COMSOL, USA) was implemented for computational simulations of relevant transport phenomena. Simulations were performed in a simplified 2D-axisymmetric geometry given the spatial symmetry of the analyzed system (**Figure S8**). To estimate the diffusion coefficient of 70 kDa dextran in the gel region, an AutoCAD file (Autodesk, USA) of the device geometry was imported and the model solved the diffusion equation:

$$\frac{\partial C}{\partial t} = D\frac{\partial^2 C}{\partial x^2} \quad (8)$$

where C is the molar concentration of the diluted molecule and *D* is the diffusion coefficient. Initial, boundary conditions of normalized concentrations $C_{max} = 1$ and $C_{min} = 0$ were applied at the left and right media channels, respectively. Zero flux conditions were imposed on the walls of the device. The diffusion coefficient was adjusted to match the experimental concentration profile at 2 hrs.

In an alternate model to investigate lymphatic drainage of interstitial solutes, a 2D-axisymmetric geometry was developed based on a lymphatic sprout within the gel region. The ascribed geometry of the sprout was based on the morphological characterization conducted in this study, while the gel domain extended to the full width of the center channel and the height was based on the approximated distance between sprouts (~65 µm). An additional domain (0.5 µm thick) was implemented between the sprout and gel region that represented the lymphatic endothelium. To this domain, the Kedem-Katchalsky equation was imposed which governs the flux of solute (N) across a semipermeable membrane:

$$N = P_e\,\Delta C + (\Delta p_{EC}\,L_p)\cdot C\,(1-\sigma_f) \quad (9)$$

where the diffusive flux is driven by the concentration difference times the diffusive permeability, and the convective flux is given by the scalar product between the fluid velocity (obtained from the Starling equation) and the local concentration. The convective term also incorporates a filtration reflection coefficient ($\sigma_f$) which considers the fraction of solutes that permeate across the endothelium along with the fluid flux. Fluid mechanics throughout the system was governed by the Brinkmann equation within the gel region and Stokes flow in nonporous regions. The transport of solutes throughout the rest of the system was governed by Fick's Second Law incorporating both diffusive and convective transport phenomena. The pressure and concentration boundary conditions were set to the same magnitudes as implemented in the experimental drainage assay with a constant value set at the inlet and a convective outflow boundary set at the outlet. Values for the intrinsic transport properties of the system ($K$, $L_p$, $P_e$) were determined experimentally in this work. Additional parameters such as the diffusion coefficient ($D$) and reflection coefficient ($\sigma_f$) for the different fluorescent-solutes were based on previous studies from our lab. Given the high degree of similarity for the diffusion coefficient value corresponding to 70 kDa-dextran from our analysis to previous studies in our lab [1–3], we approximated the diffusion coefficients of the other fluorescent-dextrans from these studies. Additionally, from a previous study [4], a range of values for the reflection coefficient were considered in which the highest bound value ascribed to the lymphatic endothelium corresponded to that measured in blood microvascular networks given that the junctions of the lymphatics are exceedingly leakier. Transient numerical solutions were generated for a total computational time of 1000 seconds, similarly as the experimental assay (~15 min). A concentration probe was added at the end of the lymphatic sprout to measure the increase in concentration during drainage (**Figure S12**). Subsequently, we quantified the corresponding drainage rate based on the concentration measurements from the last 200 seconds of the model. Additional simulations were performed to model drainage in decellularized samples, with the only corresponding difference that the domain that represented the lymphatic endothelium was removed and all the aforementioned steps were then repeated.

To model the spatial distribution of chemotactic factors during the immune recruitment assay, we extended upon the preceding model, and incorporated an additional domain consisting of solid (impermeable) spheres embedded in the gel region which represented the migrating PBMCs. Transport equations were implemented based on our previous framework, and a reaction term was also incorporated, to account for the consumption of chemokines by the immune cells, based on similar studies [5,6] ($R = 0.1\ s^{-1} \times C$). Pressure boundary conditions were set at the same magnitude as the immune recruitment experiments to establish pathological interstitial flow. A zero-concentration boundary condition was imposed at the inlet and convective outflow boundary at the outlet. Of special interest is the local secretion of chemokines by the lymphatics which was represented in the model as a constant flux condition at the endothelial domain. Despite not having experimental characterization of the secretion rates, we do have quantitative insight on the relative increase of secreted chemokines between the TNF-α-stimulated lymphatics and the unstimulated condition from the cytokine array analysis. For the diffusivity and endothelial transport properties corresponding to the chemokines, an average value was implemented based on the range of molecular weights corresponding to the chemokines of interest (CCL21, CCL19 and CXCL12) [7]. A quasi-steady state assumption was implemented in the numerical solution. Given that the timescale for diffusive and convective phenomena is 10-100 orders of magnitude less than the timescale for cell migration across the gel region (at a speed of 1-10 μm/hr) [7,8], the spatial distribution of the chemokine reaches equilibrium prior to any cellular events. From this, we are able to obtain a generalized picture on the distribution of chemokines relative to the immune cells as they migrate through the system. A series of concentration probes were drawn across the embedded spheres, which rendered the concentration profile that the immune cells would encounter as they migrate through the gel towards the lymphatics. All parameters used for the simulations are listed in Table S2.

**Transport Phenomena Analysis of Solute Drainage by Engineered On-Chip Lymphatics**

In an effort to examine the relative drainage contribution of the lymphatics across different systems, an additional solute drainage rate assessment was conducted after decellularizing the on-chip lymphatics samples. From this assessment, we were able to take into account differences in the geometry between systems to experimentally measure if the lymphatic endothelium dampens or enhances the clearance of solutes in each system. Such results demonstrated that for the monolayer lymphatics, the addition of a lymphatic endothelium reduced drainage as represented by values falling below unity when normalizing the original drainage rates by the corresponding decellularized measurements (**Figure S8**). Conversely, for both microvascular systems, we observe values greater than unity which translates to increased drainage due to the presence of the lymphatic endothelium within the 3D vascular structures. Such data suggest that the lymphatic endothelial barrier contributes to the physiological drainage functionality in engineered 3D microvascular platforms, beyond just facilitating an increased vascular surface and additional microchannels for fluid and solute transport.

Given that our previous assessment is limited to a bulk measurement of interstitial solute drainage, we sought to find an alternate study to characterize the spatial and temporal distribution of solutes during lymphatic collection and drainage. For this, we implemented an *in silico* model, developed using COMSOL Multiphysics, which recapitulated the solute drainage assay for a simplified two-dimensional geometry of a single lymphatic vessel within the gel region (**Figure S9**). From these simulations, we identified two major transport events that determined the overall flux of interstitial solutes into the lymphatics. First, as solutes enter the lymphatic vessel, driven by fluid pressure and concentration differences, the intravascular fluid carries solutes at higher flow velocities due to significantly lower hydraulic resistance imposed by the lymphatic lumen, compared to the ECM-gel region (**Figure S10**). Hence, regardless if the lumen compartment in coated by lymphatic cells or bare, solutes are preferentially transported through these microchannels, at higher rates than ECM gel systems devoid of lymphatic-like vessels. We further validated this observation using scaling arguments (see "*Scaling Analysis on Lymphatic Solute Drainage*" section) by which we confirm that the hydraulic resistance of the lumen is 3 orders of magnitude less than the resistance by the gel region. Interestingly, as solutes travel faster within these channels a second phenomenon arises where significant concentration differences develop in the transverse direction of flow. This translates into a diffusive flux of molecules that exit the lumen (which we termed as diffusive leakage) that decreases the effective amount of solutes drained by the lymphatics. While both the lymphatic and decellularized model facilitate the entry and collection of interstitial molecules to similar degrees, (**Figure S11**) the presence of a lymphatic endothelium dampens the magnitude of diffusive leakage. Thus, providing a thin barrier that prevents the diffusive-driven exit of intravascular solutes during drainage, which in turn increases the drainage rate in the lymphatic model.

To further verify the relevance of such analysis to our tissue engineered system, we extracted an additional set of data from the simulations in which a concentration probe was added at the end of the lumen channel to measure the increase in concentration during drainage, which similarly corresponds to the increase in fluorescence in our experimental system. Since the implemented model solely captures the transport of a single lymphatic sprout, direct measurements cannot be taken with our tissue-scale, experimental measurements for drainage rates. However, the ratio of drainage rates between the lymphatic sprout and the decellularize system can be applied, and taken as comparative basis to the normalized measurement that we also implemented in our experimental drainage assay. By this comparison, we found a high degree of agreement between our experimental measurements and computational results, across the different systems and for solutes/tracers of varying molecular weight (**Figure S8**); hence supporting our computational framework to describe the underlying transport phenomena and parameters that give rise to the distinctive drainage rates accordingly to the implemented system.

To further elucidate the corresponding differences in transport phenomena, we implemented scaling analysis with an emphasis on the relative timescales of solute transport within the lumen region by evaluating the Peclet number with appropriate adjustments to the scaling arguments (see "*Scaling Analysis on Lymphatic Solute Drainage*" section). We calculated that the Peclet numbers are in the range of 0.02 to 0.06, for the decellularized system, and 1.2 to 7, for the lymphatic sprout model, which implies that the diffusive timescale is significantly slower in the lymphatic vessel which hinders the leakage of intravascular solutes (**Figure S13**). Thus, enhancing the solute drainage rates compared to bare, empty channels which is also consistent with the normalized drainage measurements presented. We view this analysis as an extension on the fundamental work by Thompson et al. [9], where they study the design principles that recapitulate lymphatic drainage by pre-pattering single vessel channels followed by lymphatic cell seeding; however, their experimental study was limited to a single lymphatic-like capillary system. In this work, we are able to confirm that this biological transport phenomena also applies to a tissue-scale engineered lymphatic system.

**Scaling Analysis on Lymphatic Solute Drainage**

Numerical results depict the uniform movement of solutes across the gel region as it approaches the lymphatic sprout. Once the solutes are located at the front end of the sprout, the pressure difference across the endothelium drives the entrance of solutes into the lumen compartment. It is at this region that the transport of solutes is accelerated, compared to the solutes that continue traveling within the gel region (**Figure S10**). This local increase in transport rate can be attributed to a lower hydraulic resistance exhibited by the lumen compartment ($R_{lumen}$), as compared to the resistance imposed by the gel region ($R_{gel}$) which can be validated on the basis of scaling (Equation S1):

$$\frac{R_{gel}}{R_{lumen}} \sim \frac{\mu L / K r^2}{\mu L / r^4} \sim \frac{r^2}{K} \qquad \text{S1}$$

where the same geometric parameters, length ($L$) and radius ($r$), are attributed to each region for direct comparison, and with the dynamic viscosity ($\mu$) and hydraulic permeability ($K$) also contributing to this estimation. Upon taking the ratio of resistances, we find that the scaling analysis reduces to a comparison in length scale where the hydraulic permeability is on the order of $10^{-14}$, and the squared length of the radius is approximately $10^{-10}$ which results in a difference of 4 orders of magnitude. Thus, the lumen compartment provides a path of significantly less resistance where the solutes are preferentially transported, along with the fluid flow direction. Additionally, we can continue this analysis to verify the resistance contributed by lymphatic endothelium as Equation S2:

$$\frac{R_{endothelium}}{R_{lumen}} \sim \frac{1 / L_p \, r \, L}{\mu L / r^4} \sim \frac{r^3}{L_p \, \mu \, L^2} \qquad \text{S2}$$

where the hydraulic conductivity ($L_p$) and the surface area of the sprout provide an estimate to the resistance to fluid passage through the endothelium. For different parameters corresponding to either the growth factor- or high flow-grown lymphatics, this ratio results in a value of either 0.7 or 0.3, respectively, which suggests that the endothelium does not act as a substantial barrier to fluid transport. Since the resistance by the lymphatic endothelium is comparable to that of the lumen, then both are negligible compared to the resistance imposed by the gel region. To further validate this, we also compared the hydraulic resistance between the gel and endothelium as Equation S3:

$$\frac{R_{gel}}{R_{endothelium}} \sim \frac{\mu L / K r^2}{1 / L_p \, r \, L} \sim \frac{L_p \, \mu \, L^2}{K \, r} \qquad \text{S3}$$

from which we obtain a difference of at least 3 orders of magnitude, thus affirming that both resistances contributed by the lymphatic sprout (endothelial and luminal) are exceedingly lower than transport across the gel. Overall, these scaling arguments reveal that lymphatic sprouts facilitate solute drainage by providing a faster pathway for solute convection with minimal hindrance to fluid transport. This aligns with our previous experimental observations that 3D lymphatics achieve higher solute drainage rates, compared to a monolayer system.

To understand the underlying differences in transport phenomena between the lymphatic sprout and decellularized system, we further implemented scaling analysis with an emphasis on the relative timescales of solute transport within the lumen region. For this, we evaluated the Peclet number with appropriate adjustments to the scaling arguments accordingly to the studied system (Equation S4). For the lymphatic sprout model, the scaling parameters for the Peclet number follow as:

$$Pe_{lymph,\parallel} = \frac{(1-\sigma_f)u/L}{D/L^2} = \frac{(1-\sigma_f)u\,L}{D} \qquad \text{S4}$$

where $u$ indicates the average luminal velocity of the fluid, $L$ corresponds to the sprout length, $D$ continues to indicate the diffusion coefficient of the molecule and the reflection coefficient ($\sigma_f$) corrects for hindrance effects on the solutes. As we noted earlier, the timescale approximations in this analysis considers the relative competition of each phenomenon in the same, parallel direction. On a similar basis, the Peclet number for the decellularized sprout would be described as Equation S5:

$$Pe_{decell,\parallel} = \frac{u/L}{D/L^2} = \frac{u\,L}{D} \qquad \text{S5}$$

which simplifies to the traditional Peclet number expression. For both of these parameters, the range of values come out to be from 15 to 47 for the lymphatic sprout model, and from 24 to 77 for the decellularized system. Thus, the convective flux of solutes dominates their luminal transport, for both systems, which is in line with our previous analysis that this fluid pathway provides a faster route for solute drainage. However, diffusion of solutes simultaneously occurs at the lateral/radial direction which is responsible for the solute leakage from the lumen into the gel region observed in our computational results. As such, the Peclet number for this analysis (Equation S6) would consider the diffusive transport timescale in the radial direction as:

$$Pe_{decell,\perp} = \frac{u/_L}{D/_{r^2}} = \frac{u\,r^2}{D\,L} \qquad \text{S6}$$

for the decellularized system. Similarly, modifying the scaling arguments (Equation S7) for the diffusive rate in the lymphatic sprout yields:

$$Pe_{lymph,\perp} = \frac{(1-\sigma_f)^{u}/_L}{P/_r} = \frac{(1-\sigma_f)\,u\,r}{P\,L} \qquad \text{S7}$$

where $P$ is the diffusive permeability of the endothelium. Taking this new parameterization for the relative transport rate, we calculated that the Peclet numbers are in the range of 0.02 to 0.06, for the decellularized system, and 1.2 to 7, for the lymphatic sprout model, which implies that the diffusive rate by which these solutes are leaking out of the lumen has a significant contribution in the overall drainage in both model systems. However, the presence of the lymphatic endothelium dampens the relative magnitude of this diffusive leakage, as demonstrated in the simulation results and scaling analysis. Thus, enhancing the solute drainage rates compared to bare, empty channels which is also consistent with the normalized drainage measurements presented in the previous experimental section.

### Supplementary Tables

**Table S1: Experimental and computational default parameters for lymphatic drainage model.**

| Parameter | Symbol | Value |
|---|---|---|
| Source concentration of 3 kDa dextran | $c_{o,\ 3kDa}$ | 3.3 x $10^{-2}$ mol/m$^3$ |
| Source concentration of 10 kDa dextran | $c_{o,\ 10kDa}$ | 1.0 x $10^{-3}$ mol/m$^3$ |
| Source concentration of 70 kDa dextran | $c_{o,\ 70kDa}$ | 1.4 x $10^{-3}$ mol/m$^3$ |
| Diffusion coefficient of 3kDa dextran | $D_{3kDa}$ | 14.5 x $10^{-11}$ m$^2$/s |
| Diffusion coefficient of 10 kDa dextran | $D_{10kDa}$ | 9 x $10^{-11}$ m$^2$/s |
| Diffusion coefficient of 70 kDa dextran | $D_{70kDa}$ | 4.5 x $10^{-11}$ m$^2$/s |
| Hydraulic conductivity of monolayer lymphatics | $L_{p\ M}$ | 3.9 x $10^{-6}$ m/Pa s |
| 3 kDa dextran permeability in monolayer lymphatics | $P_{M,\ 3kDa}$ | 7.0 x $10^{-8}$ m/s |
| 10 kDa dextran permeability in monolayer lymphatics | $P_{M,\ 10kDa}$ | 1.3 x $10^{-8}$ m/s |
| 70 kDa dextran permeability in monolayer lymphatics | $P_{M,\ 70kDa}$ | 1.2 x $10^{-8}$ m/s |
| Hydraulic conductivity of growth factor-grown lymphatics | $L_{p\ GF}$ | 3.6 x $10^{-6}$ m/Pa s |
| 3 kDa dextran permeability in growth factor-grown lymphatics | $P_{GF,\ 3kDa}$ | 2.3 x $10^{-8}$ m/s |
| 10 kDa dextran permeability in growth factor-grown lymphatics | $P_{GF,\ 10kDa}$ | 4.3 x $10^{-9}$ m/s |
| 70 kDa dextran permeability in growth factor-grown lymphatics | $P_{GF,\ 70kDa}$ | 3.8 x $10^{-9}$ m/s |
| Hydraulic conductivity of high flow-grown lymphatics | $L_{p\ HF}$ | 8.1 x $10^{-6}$ m/Pa s |
| 3 kDa dextran permeability in high flow-grown lymphatics | $P_{HF,\ 3kDa}$ | 7.2 x $10^{-8}$ m/s |
| 10 kDa dextran permeability in high flow-grown lymphatics | $P_{HF,\ 10kDa}$ | 3.0 x $10^{-8}$ m/s |
| 70 kDa dextran permeability in high flow-grown lymphatics | $P_{HF,\ 70kDa}$ | 2.4 x $10^{-8}$ m/s |
| Reflection coefficient of 3 kDa dextran | $\sigma_{f,\ 3kDa}$ | 0.2-0.4 |
| Reflection coefficient of 10 kDa dextran | $\sigma_{f,\ 10kDa}$ | 0.4-0.8 |
| Reflection coefficient of 70 kDa dextran | $\sigma_{f,\ 70kDa}$ | 0.4-0.8 |

**Table 2: Experimental and computational default parameters for chemokine transport model.**

| | | |
|---|---|---|
| Diffusion coefficient of chemokines | $D_{ch}$ | $14.5 \times 10^{-11}$ m$^2$/s |
| Flux of chemokines from the lymphatics | $N_{ch}$ | $1 \times 10^{-6} – 2.5 \times 10^{-6}$ mol/m$^2$ s |
| Hydraulic conductivity of high flow-grown lymphatics | $L_{p\ HF}$ | $8.1 \times 10^{-6}$ m/Pa s |
| Chemokine permeability in high flow-grown lymphatics | $P_{HF,\ ch}$ | $7 \times 10^{-8}$ m/s |
| Reflection coefficient of chemokines | $\sigma_{f,\ ch}$ | 0.2 |
| Consumption rate of chemokines by PBMCs | $R_{ch}$ | 0.5 x C mol/s |

### Supplementary Figures

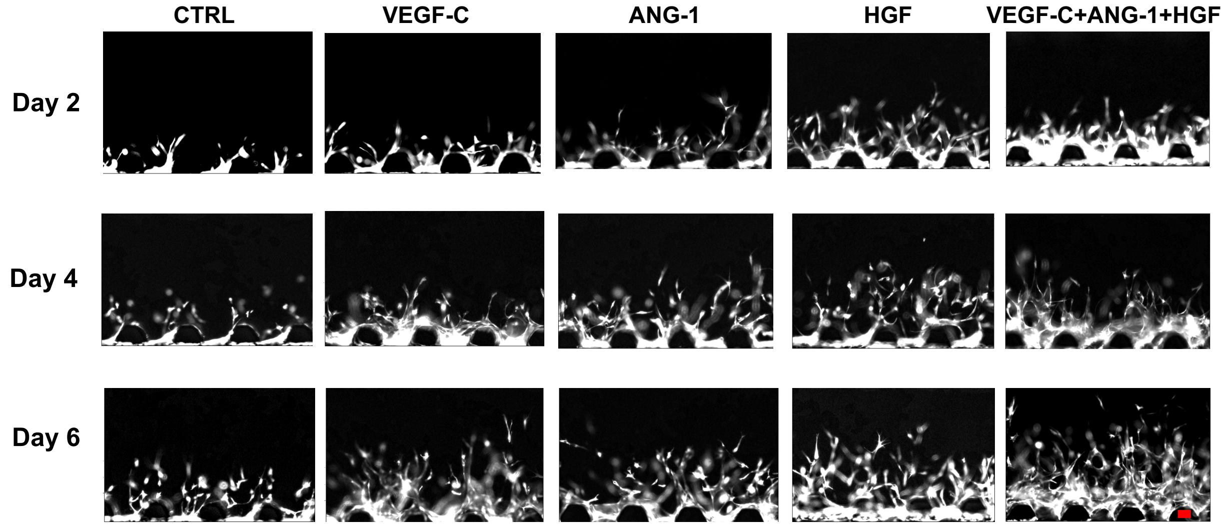


**Figure S1.** Representative images of lymphatic sprouting for different experimental conditions under biochemical stimulus with growth factors at different days. Scale bar is 100 μm.

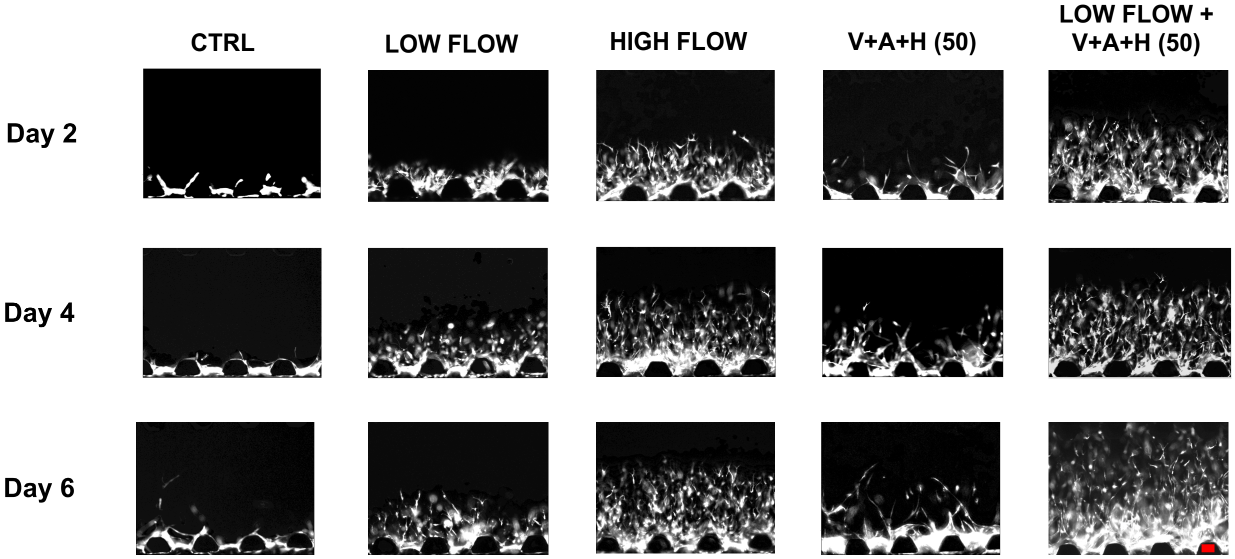


**Figure S2.** Representative images of lymphatic sprouting for different experimental conditions corresponding to stimulus by interstitial flow at different velocities. Scale bar is 100 μm.

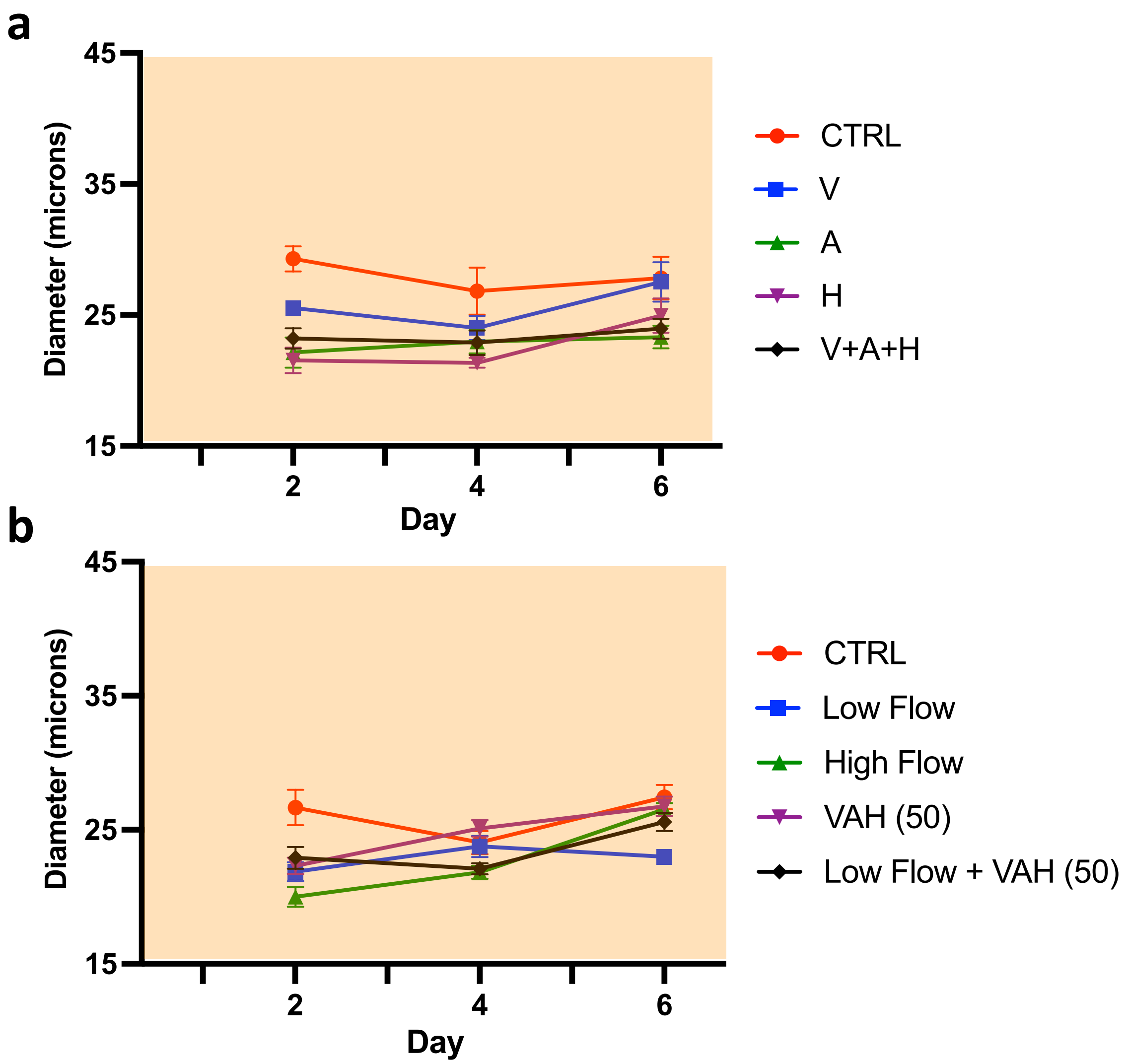


**Figure S3.** Quantitative analysis of lymphatic vessel diameters for (a) biochemical-stimulated lymphatics with growth factors and (b)interstitial flow-stimulated lymphatics. Highlighted regions correspond to *in vivo* values.

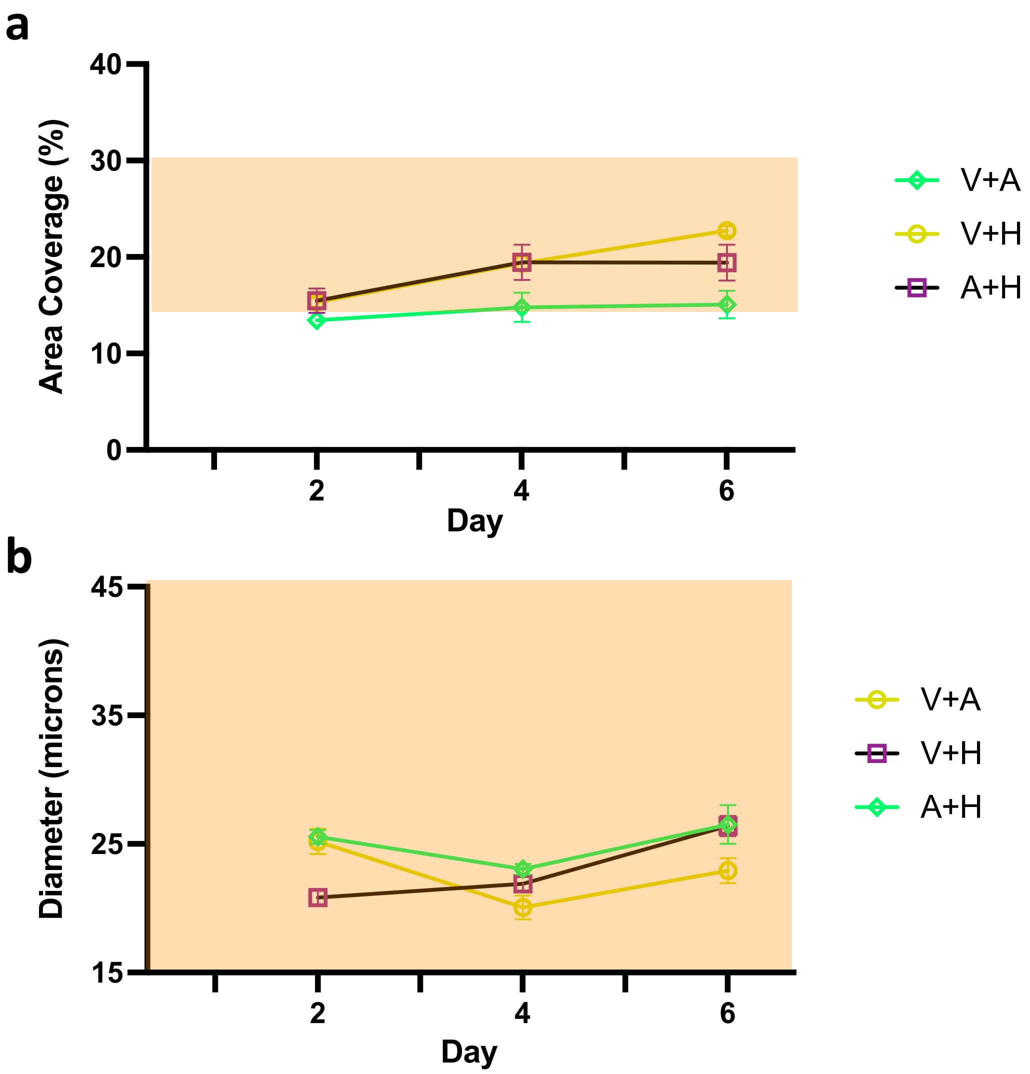


**Figure S4.** Quantitative analysis of lymphatic vessel morphology: (a) lymphatic area of coverage and (b) lymphatic vessel diameter for additional combinations of growth factors. Highlighted regions correspond to *in vivo* values.

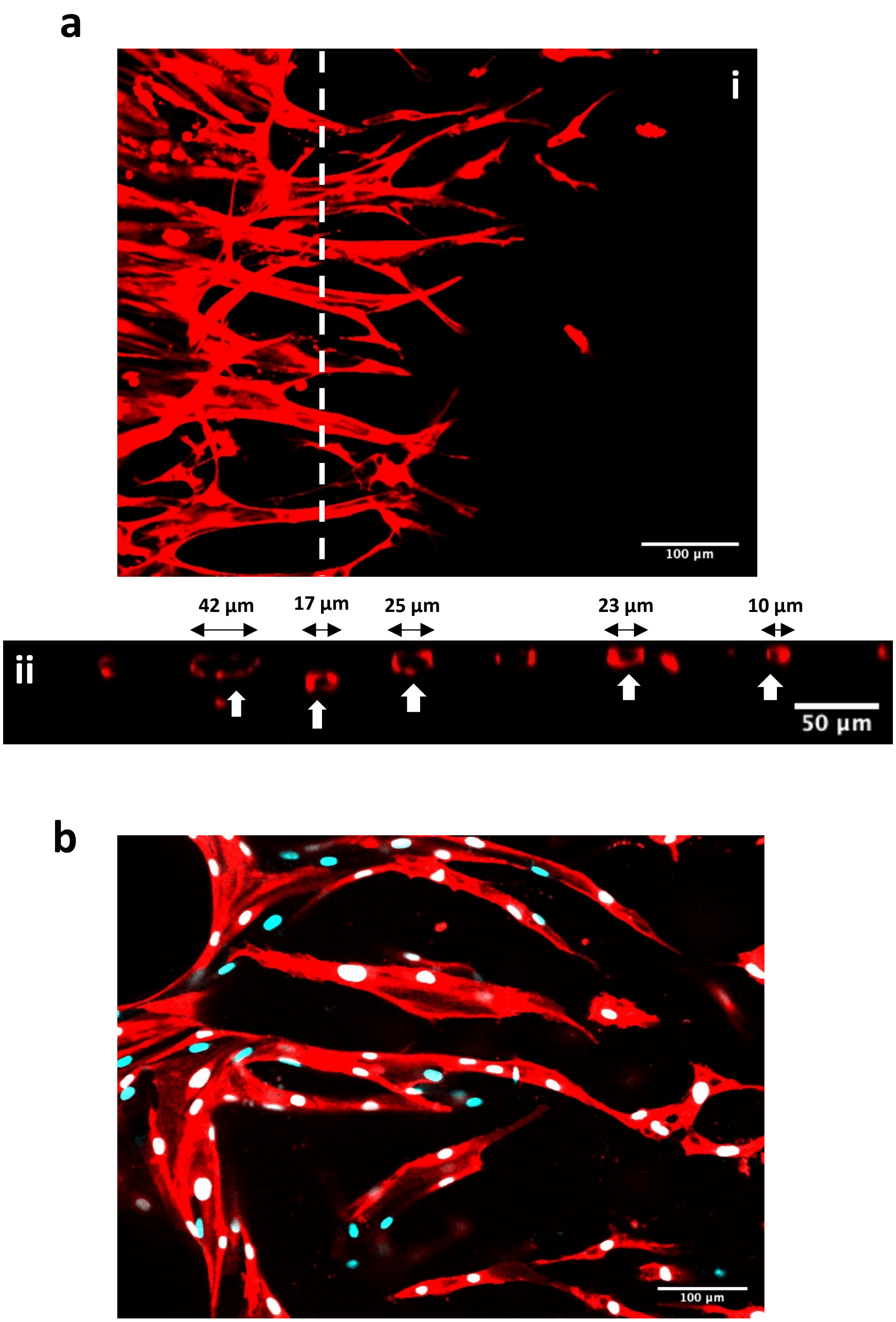


**Figure S5.** (a) Representative images of the lymphatic vasculature expressing RFP (i) with an orthogonal view (ii) of the vessel depicting lumen compartments. White arrows indicate vascular lumens. (b) Representative image of engineered lymphatic vasculature expressing RFP and stained for PROX-1 transcriptional factor (cyan). Scale bars are specified within the image.

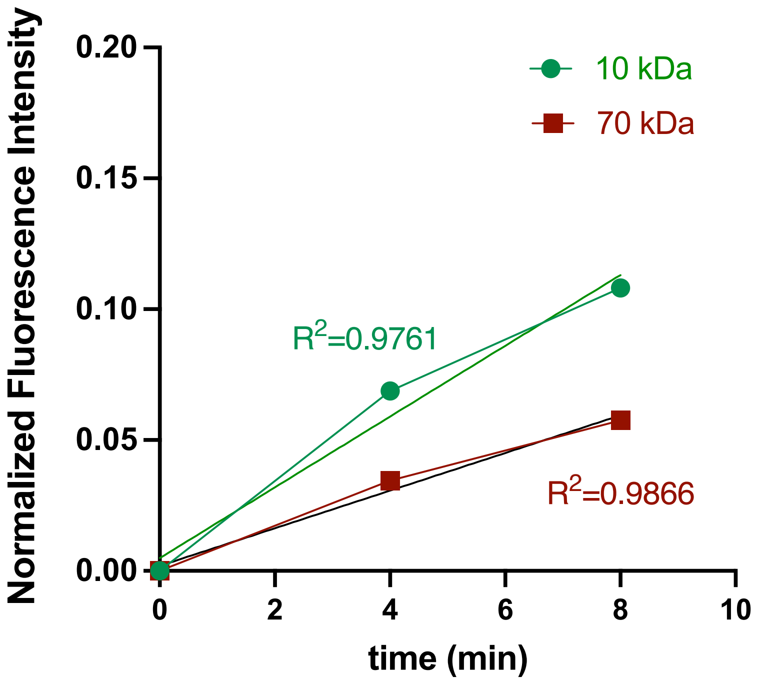


**Figure S6.** Temporal plot of fluorescence intensity corresponding to solute drainage measurements of 10 kDa dextran for high-flow engineered lymphatics.

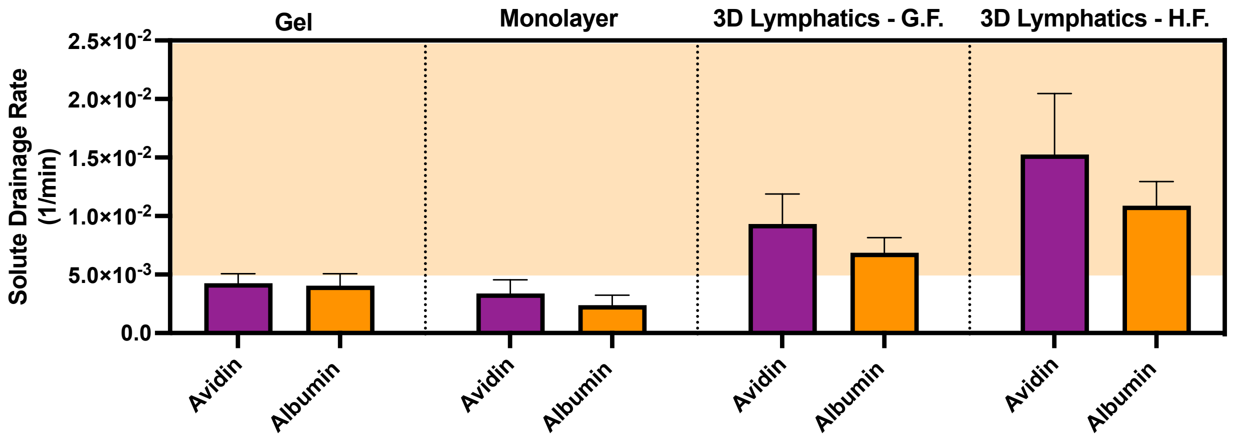


**Figure S7.** Quantification results of solute drainage rates accordingly to the experimental condition and dextrans of varying biological molecules.

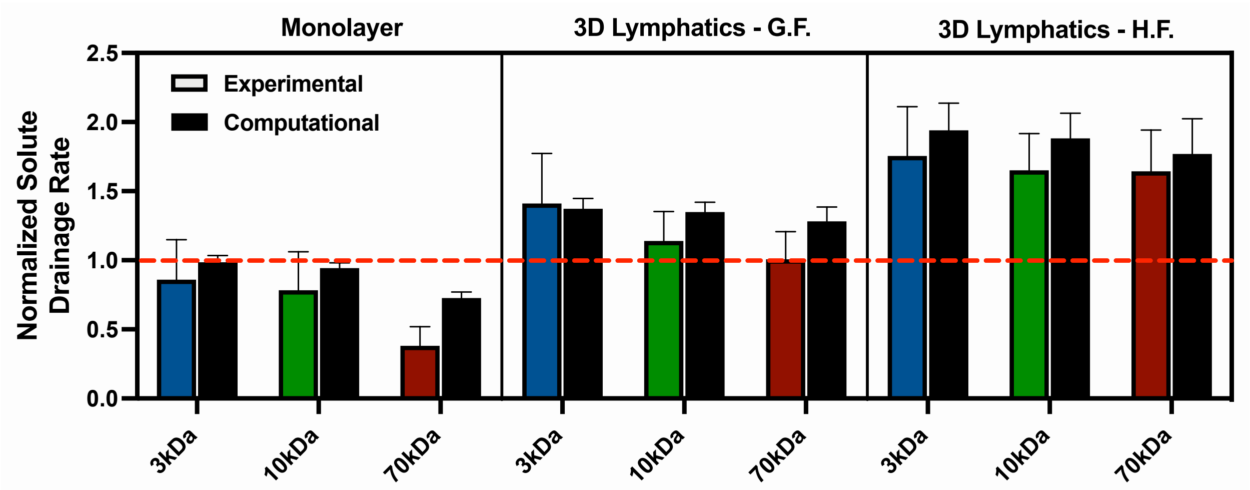


**Figure S8.** Normalized solute drainage rates (relative to the respective decellularized sample) for dextrans of varying molecular weight accordingly to the experimental condition that the lymphatics were implemented in the microfluidic system.

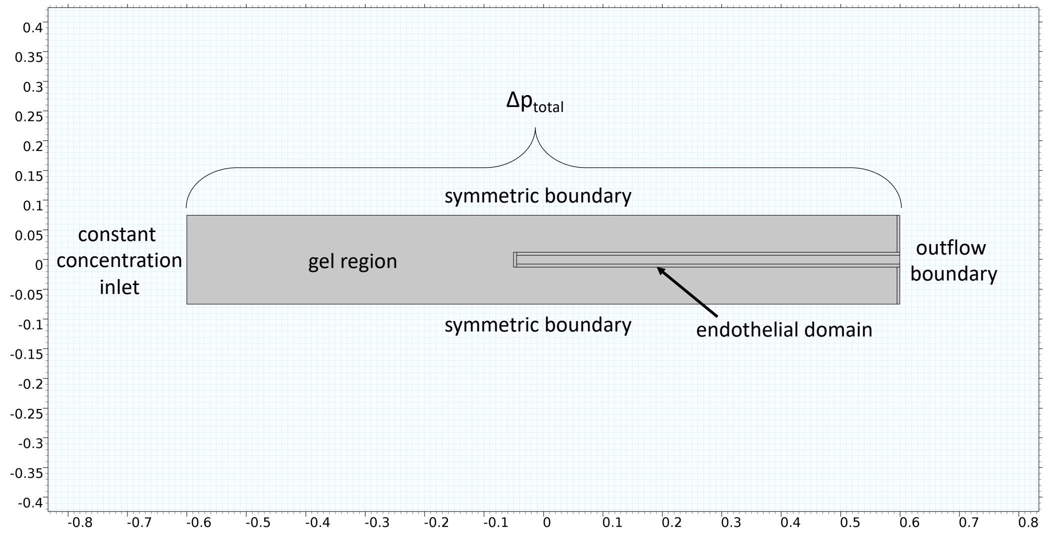


**Figure S9.** Geometry and boundary conditions of the computational model for lymphatic solute drainage developed in COMSOL Multiphysics.

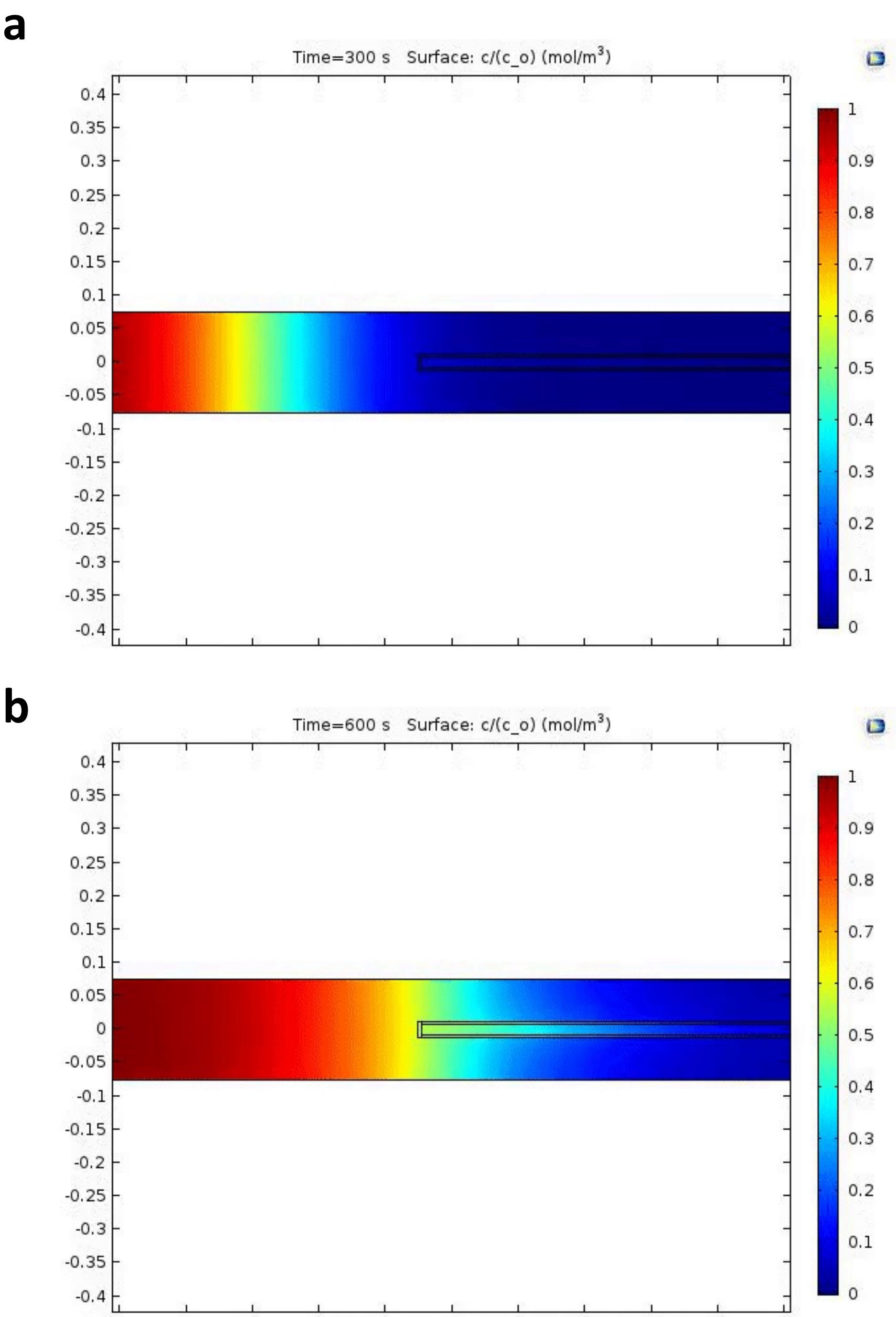


**Figure S10.** Computational model results for lymphatic solute drainage developed in COMSOL Multiphysics depicting the normalized concentration of solutes representing 10 kDa dextran at different stages during drainage: the first (upper) image showing the approaching solutes to the sprout at 300 seconds and the second (lower) image showing the spatial distribution of solutes at 600 seconds.

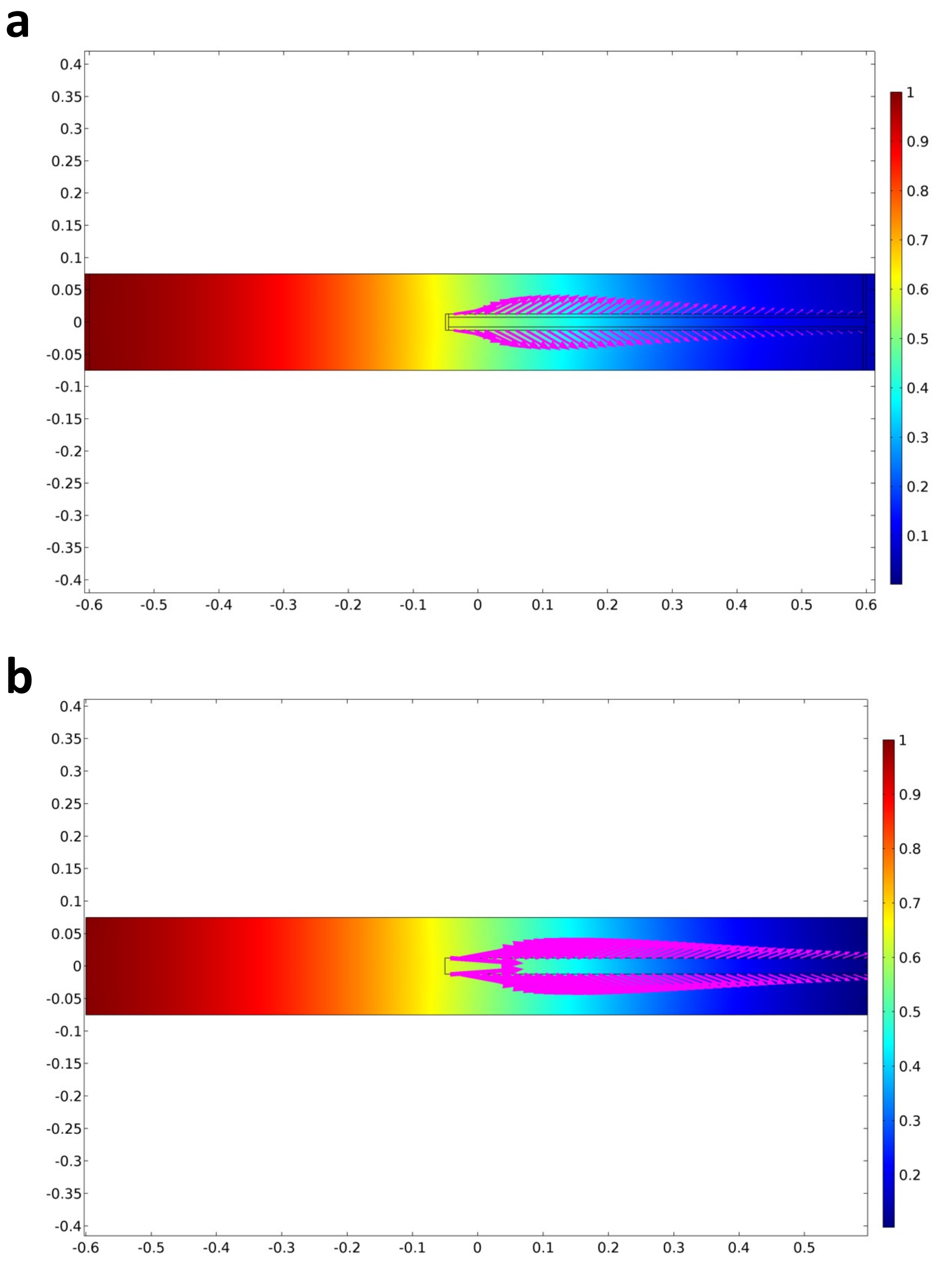


**Figure S11.** Computational model results for lymphatic sprout solute drainage (upper image) and decellularized-sprout solute drainage (lower image). Both models depict the normalized concentration of solutes representing 3 kDa at 500 seconds and the magenta-color arrows indicate the magnitude and directionality of the diffusive flux at the lumen surface.

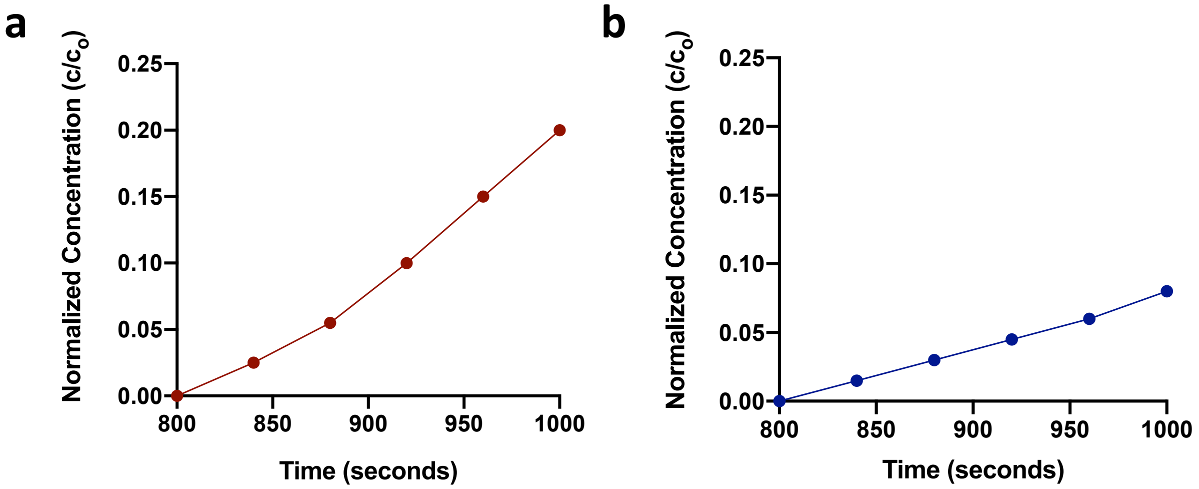


**Figure S12:** Concentration plot corresponding to a concentration probe placed at the end of the lumen compartment, thus measuring the increase in solute concentration during drainage by the lymphatic sprout model (a) and the decellularized-sprout system (b). Both plots pertain to 10 kDa dextran-based solute properties.

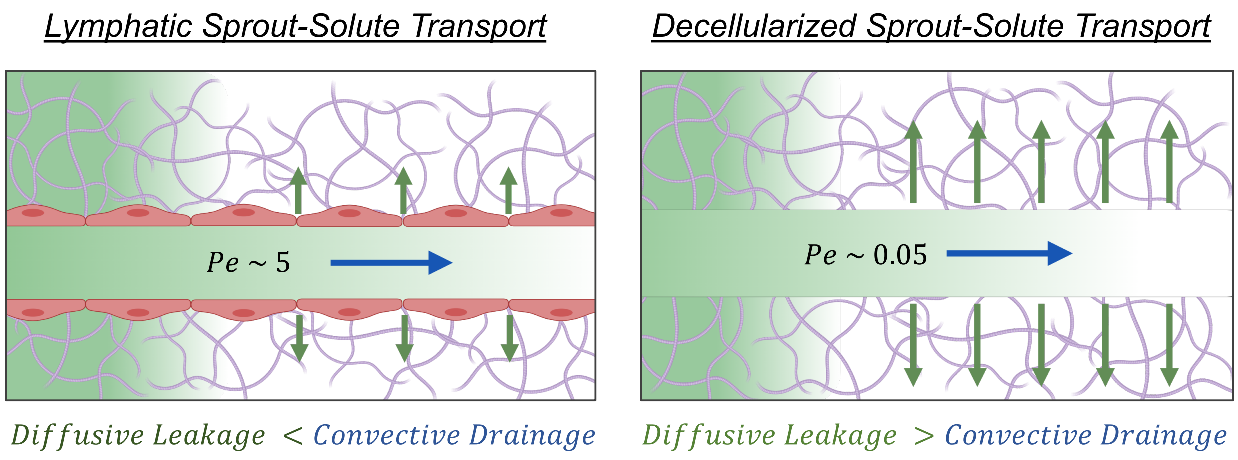


**Figure S13.** Schematic on solute transport models for lymphatic or decellularized sprouts where the relative competition between convective drainage and diffusive leakage are highlighted schematically and quantified by the Peclet number (*Pe*) correspondingly for each system. Both experimental and computational results validate the increase drainage of interstitial molecules with the presence of a lymphatic endothelium.

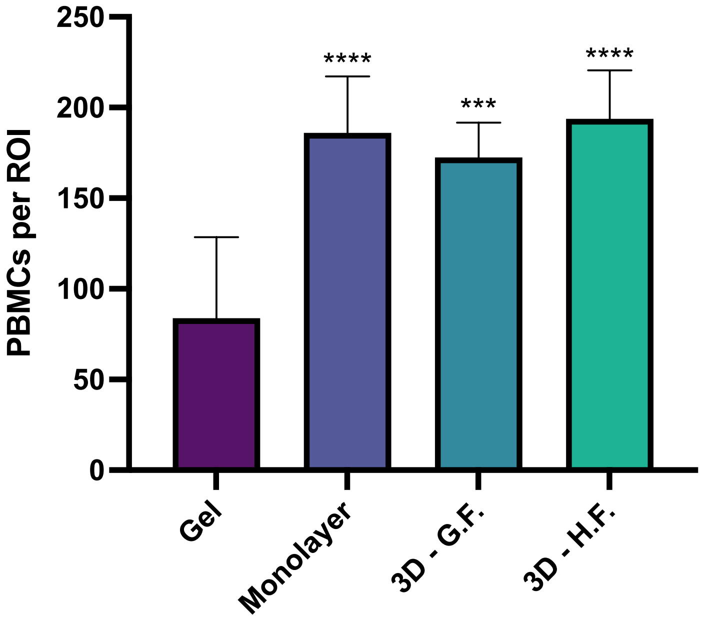


**Figure S14.** Quantitative analysis from the PBMCs infiltration assay performed on varying experimental conditions as described in the plot. Statistical significance is reported with respect to the gel sample.

| | L | C | P |
|---|---|---|---|
| **IgG** | 0.64 | 0.41 | 0.46 |
| **CCR7** | 0.05 | 0.06 | 0.51 |
| **CXCR4** | 0.02 | 0.01 | 0.70 |
| **CCR7 + CXCR4** | 0.03 | 0.12 | 0.58 |

**Figure S15.** Heat map of PBMC distribution within the gel region for specified antibody-neutralization conditions by each row, and locations by column. The table columns for L, C and P correspond to the gel region with lymphatics, central gel region and region where the PBMCs are introduced, respectively.
